# Disruption of the SYNGAP1 PDZ ligand motif accelerates differentiation of human iPSC-derived GABAergic neurons

**DOI:** 10.64898/2026.02.24.707848

**Authors:** Jianzhi Jiang, Ruslan Rust, Ilse Flores, Yaowen Feng, Peyman Nouri, Veronica A. Clementel, Amir Arya, Ali Basirattalab, Ivy Y. Yang, Antigoni Manousopoulou, Spiros D. Garbis, Nicholas A. Graham, Marcelo P. Coba

## Abstract

SYNGAP1 haploinsufficiency is a leading genetic cause of neurodevelopmental disorders (NDD), including intellectual disability and epileptic encephalopathy. While most studies on SYNGAP1 function have focused on glutamatergic neurons, its role in GABAergic neurons and during early neuronal development is unclear. Using human iPSC-derived GABAergic neurons, we demonstrate that SYNGAP1 haploinsufficiency accelerates neuronal maturation, characterized by increased dendritic length, synaptic density, and maturation of synaptic structures. Disruption of the isoform-specific SYNGAP1 PDZ binding motif reproduces these phenotypes, highlighting the critical role of PDZ-mediated interactions in regulating GABAergic neuronal differentiation. Proteomic and phosphoproteomic analyses reveal significant dysregulation of synaptic proteins, RNA processing, and transcriptional control, with a significant increase in postsynaptic density proteins content. RNA-seq analysis suggest that the acceleration in neuronal differentiation starts few hours after neuronal induction setting a path to a faster neuronal and synapse maturation. These findings establish that SYNGAP1 acts as a key regulator of neuronal differentiation across both excitatory and inhibitory neurons. Our work underscores the importance of the SYNGAP1 PDZ ligand motif for normal neuronal development and suggests translational strategies targeting SYNGAP1 alpha1 isoform levels to mitigate SYNGAP1-related NDD.

## Introduction

SYNGAP1 is one of the most prevalent genes associated with neurodevelopmental disease (NDD). Nonsense mutations leading to haploinsufficiency or reduced SYNGAP1 protein levels are a common cause of sporadic and non-syndromic intellectual disability (ID)^1–5^. Several variants of unknown significance have also been reported, and they are spread throughout the gene and are likely to affect different functional domains of the SYNGAP1 protein^2,6^. Many of these variants have been associated to autism spectrum disorders (ASD), and epileptic encephalopathy^4–11^.

SYNGAP1 function has largely been studied within the context of mature rodent excitatory synapses from glutamatergic neurons^12–18^. However, different lines of evidence suggest a role of SYNGAP1 at early stages of neuronal development^19–21^. In human induced glutamatergic neurons, the lack of SYNGAP1 protein accelerates neuronal maturation, with increases in dendritic length, number of synapses and accelerated expression of synaptic and network activity^21^. In human radial glia cells (hRGCs) a decrease in SYNGAP1 protein levels or impairments in its RASGAP activity disrupt function and accelerate the maturation of cortical projection neurons^20^. Most of our knowledge of SYNGAP1 function comes from the study of glutamatergic neurons; however, in mature rodent GABAergic neurons, it has been proposed that Syngap1 haploinsufficiency impairs terminal axonal branching and bouton formation of cortical parvalbumin positive neurons^22^. However, it is not known if SYNGAP1 function is necessary during the early stages of neuronal differentiation in GABAergic neurons. Importantly, it is not known what functional domains in SYNGAP1 are needed for its role in inhibitory neurons.

SYNGAP1 mutation can affect different functions, including its RASGAP activity, the capacity to associate to PDZ domains or to reduce SYNGAP1 total protein levels. While a decrease in SYNGAP1 protein levels and RASGAP activity dysregulate neuronal development in human models^20^, the role of SYNGAP1 PDZ ligand function in neuronal development, is not fully known. However, the capacity of SYNGAP1 to associate to PDZ-containing proteins is isoform-dependent and exclusive of the SYNGAP1 **alpha** (𝞪)1 isoform^13,15,23,24^. This isoform contains a PDZ ligand motif that has the capacity to associate SYNGAP1 to glutamatergic postsynaptic density (PSD) protein interaction networks^13,25–28^. It has been shown that Syngap1𝞪1 functions as a negative regulator of mature excitatory synapse structure in rodent models^4,29,30^. As well, the disruption of Syngap1𝞪1 isoform impairs synaptic plasticity, and learning^13^. However, it is not known if the SYNGAP1 PDZ ligand function is necessary for normal neuronal development.

Here, we show that SYNGAP1 haploinsufficiency accelerates the maturation of GABAergic induced neurons (iNs) and that the disruption of the SYNGAP1 PDZ ligand motif is enough to disrupt the speed of differentiation of human GABAergic neurons. We characterized the proteome of developmental human iPSC-derived GABAergic iNs and show that SYNGAP1 is enriched in PSD fractions. We show that SYNGAP1 PDZ ligand dysfunction increases the length of dendritic branches, synapse density and synaptic maturation, increasing protein levels of presynaptic and postsynaptic components of GABAergic iNs. Analysis of phosphoproteome shows that the lack of capacity of SYNGAP1 to associate to PDZ containing proteins dysregulates synaptic, postsynaptic and signaling mechanisms modulating RNA processing and transcriptional control. RNA-seq analysis of iPSC SYNGAP1 mutant lines shows alterations on neuronal differentiation as early as 12hs post neuronal induction. These results suggest that the capacity of SYNGAP1 to negatively regulate neuronal differentiation is not exclusive to glutamatergic neurons and that SYNGAP1 PDZ ligand is necessary for SYNGAP1 function during early stages of human GABAergic neuronal differentiation.

## Results

To start to address the role of SYNGAP1 in developing human inhibitory neurons we first derived GABAergic induced neurons (iNs) by direct differentiation from iPSCs using a modified protocol via induced expression of ASCL1 and DLX2 as previously described^31^. First, we confirmed that GABAergic iNs expressed somatostatin (SST) and GAD1 markers (Fig. 1A). Then, to confirm that GABAergic iNs, were able to release GABA, we used the genetically encoded GABA sensor iGABASnFR^32^. We transfected GABAergic iNs with a AAV2/1.hSynapsin1.iGABASnFR virus and monitored GABA release by live imaging at 14 and 21 DIV. As control, we transfected Glutamatergic iNs, induced via expression of NGN2 as we previously described^33^. Fig.1B shows that the iGABASnFR sensor was able to respond only in GABAergic iNs with no signal above background being detected in Glutamatergic iNs.

**Figure 1.**
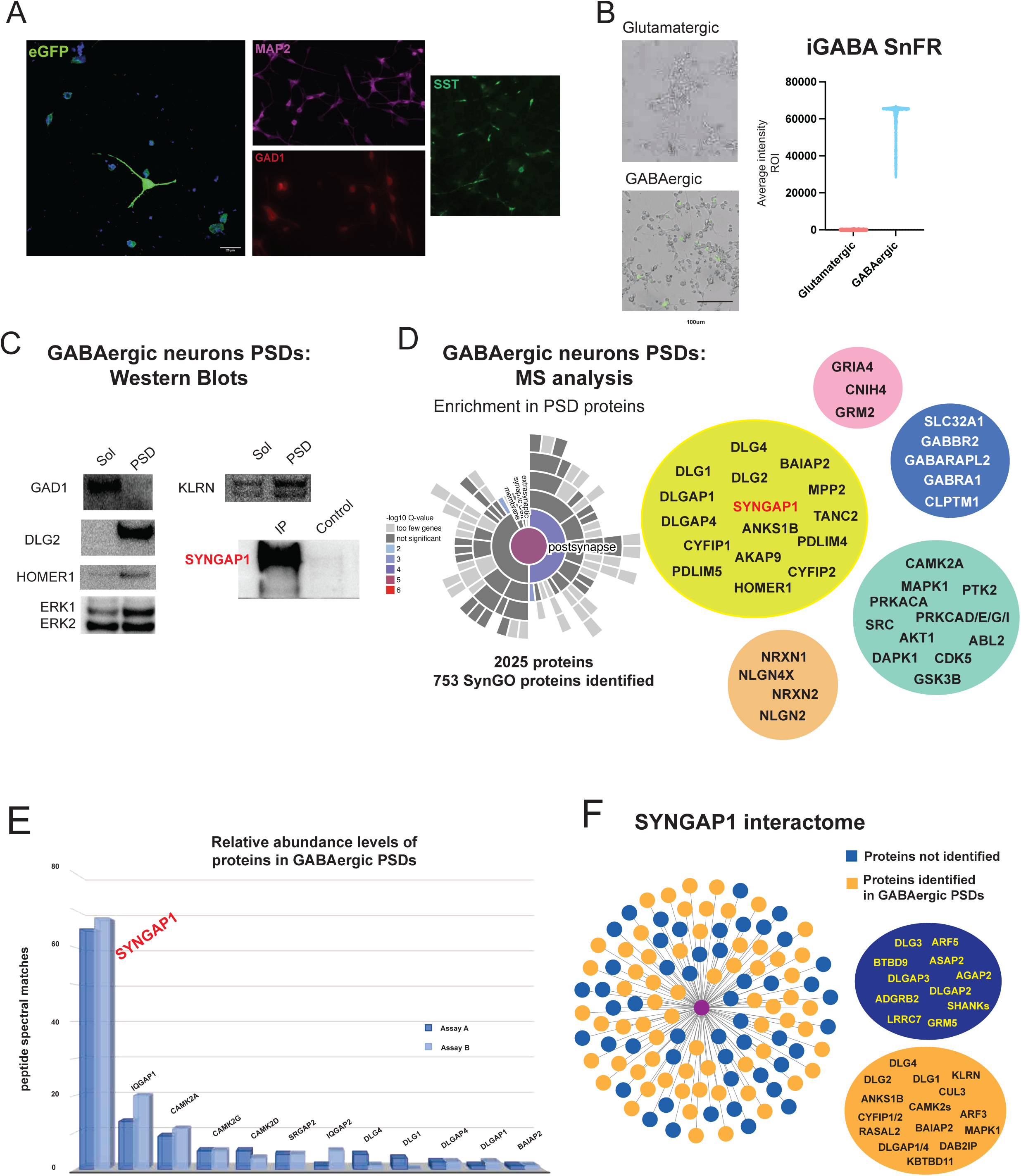
A. At 7DIV GABAergic neurons, expression of eGFP together with MAP2, GAD1 and somatostatin (SST). B. Response to the activation of the GABAergic sensor iGABA SnFR in glutamatergic (top panel) and GABAergic neurons (lower panel). Graph shows average intensity recorded from GABAergic induced neurons (iN) in response to neuronal release of GABA. No responses above background are detected in glutamatergic iN (Triplicate assays from 2 independent differentiations). C. Representative western blots (WB) from postsynaptic density (PSD) enriched fractions. Blots show comparisons between PSD enriched and PSD depleted fractions from 21DIV GABAergic iN. Blots show a pattern consistent with PSD enriched fractions with similar levels of ERK1/2, increased levels of PSD93, Homer1, Kalirin (KLRN) and no presence of GAD1 in PSD fractions. Right/bottom blot shows immunoprecipitation and WB of SYNGAP1 from PSD fractions compared against a SYNGAP1 KO cell line^20^ as control. The same set of samples used for WB (C) were analyzed by Mass Spectrometry (D). D. Sun plot shows SynGO analysis of PSD enriched fractions from GABAergic iN. From a total of 2025 proteins, 753 were identified as SynGO synaptic proteins with an enrichment in PSD proteins (postsynapse). Representative proteins from the core signaling machinery of the PSD are grouped in Yellow: scaffolds, Pink: glutamatergic signaling, Blue: GABAergic signaling, Cyan: protein kinases, and Orange: Synaptic cell adhesion. E. Bar graph showing relative abundance of core components of the PSD identified in GABAergic enriched fractions from 21 DIV iN. Bar plot shows a high relative abundance of SYNGAP1 in comparison with its known protein interactors and PSD scaffolds such as DLG4 (PSD95), DLG1, IQGAP2, DLGAP1/4 (Plot represents results from replicate assays A and B). F. Comparison between the Syngap1 protein interactome as reported in mature rodent synapses and protein interactors identified (orange) and not identified (blue) in GABAergic iN. Examples of both, identified and not identified proteins, are shown in orange and blue balloons. Red node is Syngap1.

### Molecular characterization of PSD enriched fractions from human GABAergic neurons

To characterize and determine the expression of SYNGAP1 in GABAergic iN we isolated postsynaptic density (PSD) enriched fractions using a similar protocol as we previously described for Glutamatergic iNs^33^. Their molecular composition was analyzed in duplicate assays by Western Blot (WB) and Mass Spectrometry (MS) (Fig. 1C. Suppl. Table 1). We identified 2025 unique proteins, from both datasets, with 753 proteins validated as synaptic proteins by SynGO 1.3 analysis. Although SynGO is biased to reported synaptic proteins from adult rodent synapses^34^, our analysis still shows an enrichment of postsynaptic proteins (Fig.1D. Suppl. Table 1). In contrast to induced glutamatergic neurons^33^, we were not able to identify the glutamatergic markers SLC17A7 and SLC17A6 (Suppl. Table 1). Synaptic fractions contain core scaffold proteins such as DLGs/DLGAPs/HOMERs, ionotropic and metabotropic glutamate receptors, protein kinases such CAMK2s and synaptic adhesion proteins such as NRXNs/NLGNs (Fig. 1D. Suppl. Table 1), together with GABAergic signaling components including the GABAergic neuron marker SLC32A1 (Fig. 1 Suppl. Table 1). We also identified SYNGAP1 in PSDs and in SYNGAP1 immunoisolated fractions from GABAergic iNs (Fig. 1C). SYNGAP1 is, relatively, one of the most abundant proteins in PSD enriched fractions of GABAergic iNs, with relative levels compared to cytoskeletal proteins (Suppl. Table 1). This is more evident when compared to the relative levels observed for DLGs (PSD95, PSD93, SAP97) or members of the CAMK2 family (CAMK2𝞪, beta, gamma, and delta) (Fig. 1E. Suppl Table 1).

The analysis of the SYNGAP1 interactome shows several components of GABAergic PSDs that have been also described in adult rodent synapses (Fig. 1F. Suppl. Table 1, 2)^26,27^. These include direct SYNGAP1 interactors such as PSD95, PSD93, SAP97, ANKS1B and other components of the interactome CAMK2s, BAIAP2, CYFIP1/2, DLGAP1/4 (Fig. 1D and 1F. Suppl. Table 2). A number of these proteins contain PDZ protein domains suggesting that SYNGAP1 might associate to these proteins using its PDZ-ligand interface as reported in mature rodent synapses^13,15,26–28^.

### SYNGAP1 haploinsufficiency accelerates the differentiation of GABAergic iNs

We previously described^20^ that the SYNGAP1 patient derived mutation p.Q503X (SYNGAP1 p.Q503X (patient)), causes a haploinsufficient model of SYNGAP1 dysfunction with an acceleration in the maturation of cortical projection neurons^20^. Here, we asked if the decrease in SYNGAP1 total protein levels also influences the maturation of GABAergic neurons. We first observed that a reduction in SYNGAP1 total protein levels increased the number of SST (Somatostatin) positive neurons at 4, 10 and 21 DIV, suggesting an acceleration in the development of GABAergic iNs (Fig. 2A-C). The SYNGAP1 p.Q503X (patient) iNs also have a significant increase in the length of dendrites with more spines and higher spine density. The spines were also more mature with a significant increase in the number of mushroom spines (Fig. 2D), suggesting that a decrease in total SYNGAP1 protein levels produces an acceleration of GABAergic neuronal and spine maturation.

**Figure 2.**
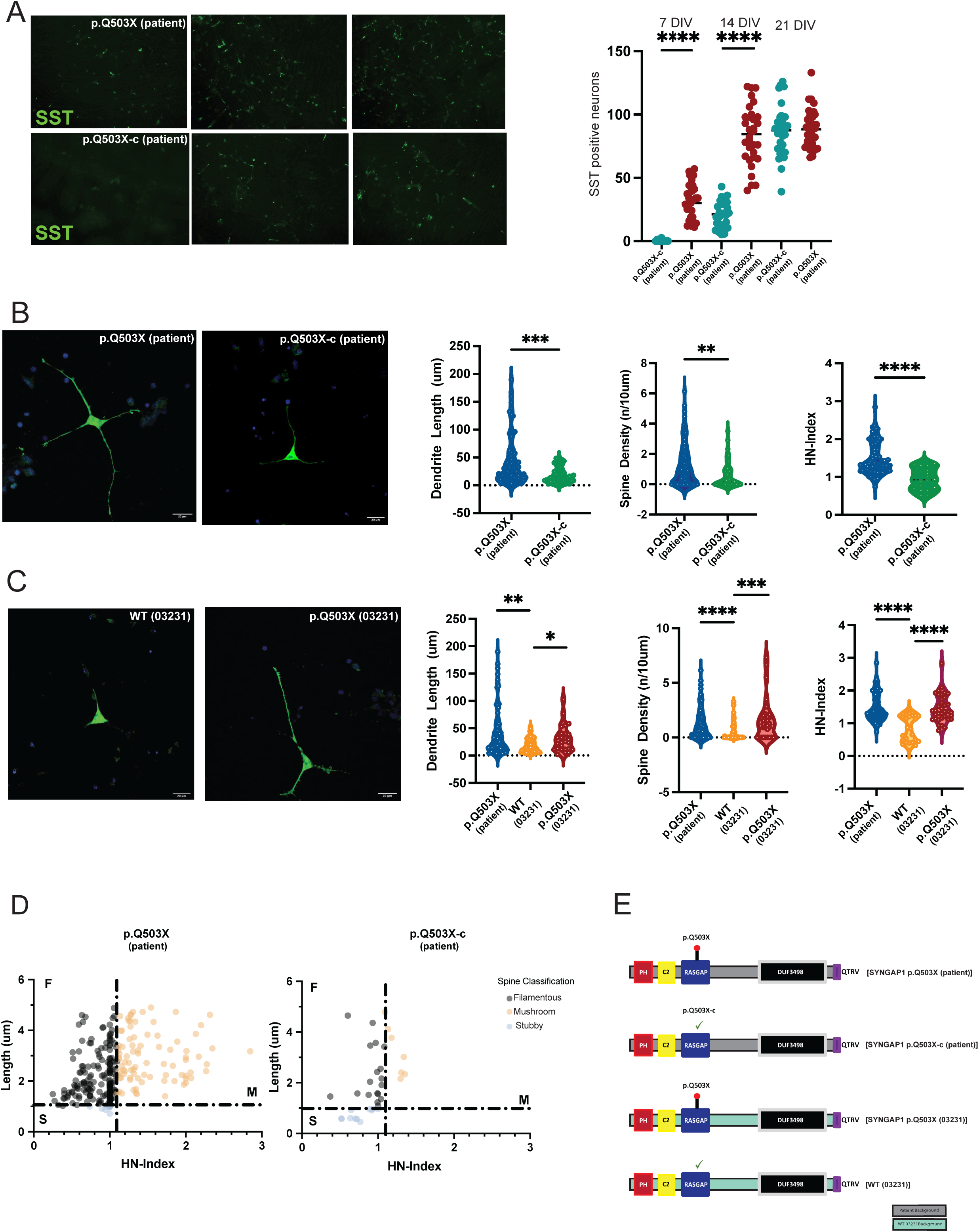
A. Identification of somatostatin (SST) positive neurons at 4, 10 and 21DIV. Graph shows a significant increase (two-tailed Student’s *t* test p<0.01) in the identification of SST positive neurons in the patient-derived SYNGAP1 haploinsufficient cell line p.Q503X at 4 and 10DIV, when compared to its isogenic control (p.Q503X-c (patient)). Results corresponds to 4 replicates from three independent differentiations. B. Representative confocal max intensity projection images of cultured GABA-iNs at 9 DIV after GFP-transfection at 6DIV from the patient derived mutation p.Q503X corrected cell line (p.Q503X-c (patient)), and the patient derived p.Q503X mutation (p.Q503X (patient)). Violin plots show quantification dendritic length, spine density, and head-neck (HN)-Index, in each cell line. The number of spines per 10um of 40-80um dendritic segments p=0.0004), dendritic length (p=0.0027) and HN index (p<0.0001) is significantly higher in SYNGAP1 haploinsufficient p.Q503X (patient) compared to its isogenic control (p.Q503X-c (patient)). C. Analysis of SYNGAP1 p.Q503X mutation introduced into the WT 03231 background p.Q503X (03231), compared to its correspondent isogenic control WT (03231). Results show a significant increase in dendritic length (p=0.0022), spine density (, p<0.0001) and HN index (p<0.0001) when compared against its isogenic control WT (03231). No significant differences are observed between the p.Q503X (patient) and p.Q503X (03231) GABAergic iN, with similar significant results obtained in both backgrounds. D. Head-neck(HN)-Index of spines was significantly higher in both p.Q503X (patient) and p.Q503X mutation inserted into the WT (03231) normal background (p.Q503X (03231)). Individual non-filopodial spines were plotted according to their length (um) and HN-Index for each line. Figure shows an increase in the number of filamentous and mushroom spines in GABAergic iN, in the SYNGAP1 model of haploinsufficiency p.Q503X (patient) and p.Q503X (03231). For each cell line, 6 replicates were plated from at least three differentiations. Two-sided Mann-Whitney testing was used. Differences between more than two groups were analyzed by one way-ANOVA with Tukey correction for multiple testing, unless the data was non-normally distributed for which a Kruskall-Wallis H test was used. Error bars represent the s.e.m. D. Distribution of filamentous, stubby and mushroom spines for p.Q503X (patient) and its correspondent isogenic control. E. Cartoon representation of the SYNGAP1 mutations and cell lines used. The cartoon shows the haploinsufficient mutation p.Q503X in the patient background (grey) and the WT (03231) background (cyan), together with the WT (03231) control background. All the cell lines have normal PDZ ligand QTRV (purple).

While all these comparisons were performed against the correspondent isogenic control (patient line with the mutation corrected): SYNGAP1 p.Q503X-c(patient), we wanted to confirm that the observed effects of the SYNGAP1 p.Q503X mutation were not due to genetic background effects. For this purpose, we introduced the SYNGAP1 p.Q503X into the “wildtype” 03231) control-iPSC line (WT(03231)) derived from a healthy 56-year-old: ((SYNGAP1 p.Q503X (03231)) as we previously reported^20^. Fig. 2C shows that SYNGAP1 p.Q503X (03231) cell line produced GABAergic iNs with the same phenotype observed for the patient-derived cell line, SYNGAP1 p.Q503X (patient). This indicates that the increase in the acceleration of neuronal and spine maturation observed is not due to genetic background effects. The different cell lines, with their corresponding backgrounds, used for the experiments are represented in Fig. 2E.

### SYNGAP1 haploinsufficiency accelerates iPSCs differentiation into GABAergic neurons

We have previously shown that in hRGCs a decrease in SYNGAP1 protein levels produces an acceleration in the differentiation of cortical projection neurons^20^. Here, we determined that as early of 7DIV, a reduction of SYNGAP1 total protein levels, produced an acceleration in the differentiation of SST positive neurons. Therefore, we wanted to determine if SYNGAP1 produced an acceleration of differentiation of iPSCs into GABAergic neurons before they became SST positive neurons. Therefore, we performed RNA-seq analysis of the SYNGAP1 haploinsufficient p.Q503X mutation and its correspondent isogenic controls at 5 different time points: T=0h (iPSC stage), and 12hs, 24hs, 48hs and 96hs post GABAergic neuron induction (Fig. 3A). Principal component analysis reveals an early separation between P1 and WT cells beginning at 12 hours of differentiation (Fig 3B). This divergence becomes progressively more pronounced at between 48 and 96 hours, indicating sustained transcriptional divergence over time (Fig. 3B).

**Figure 3.**
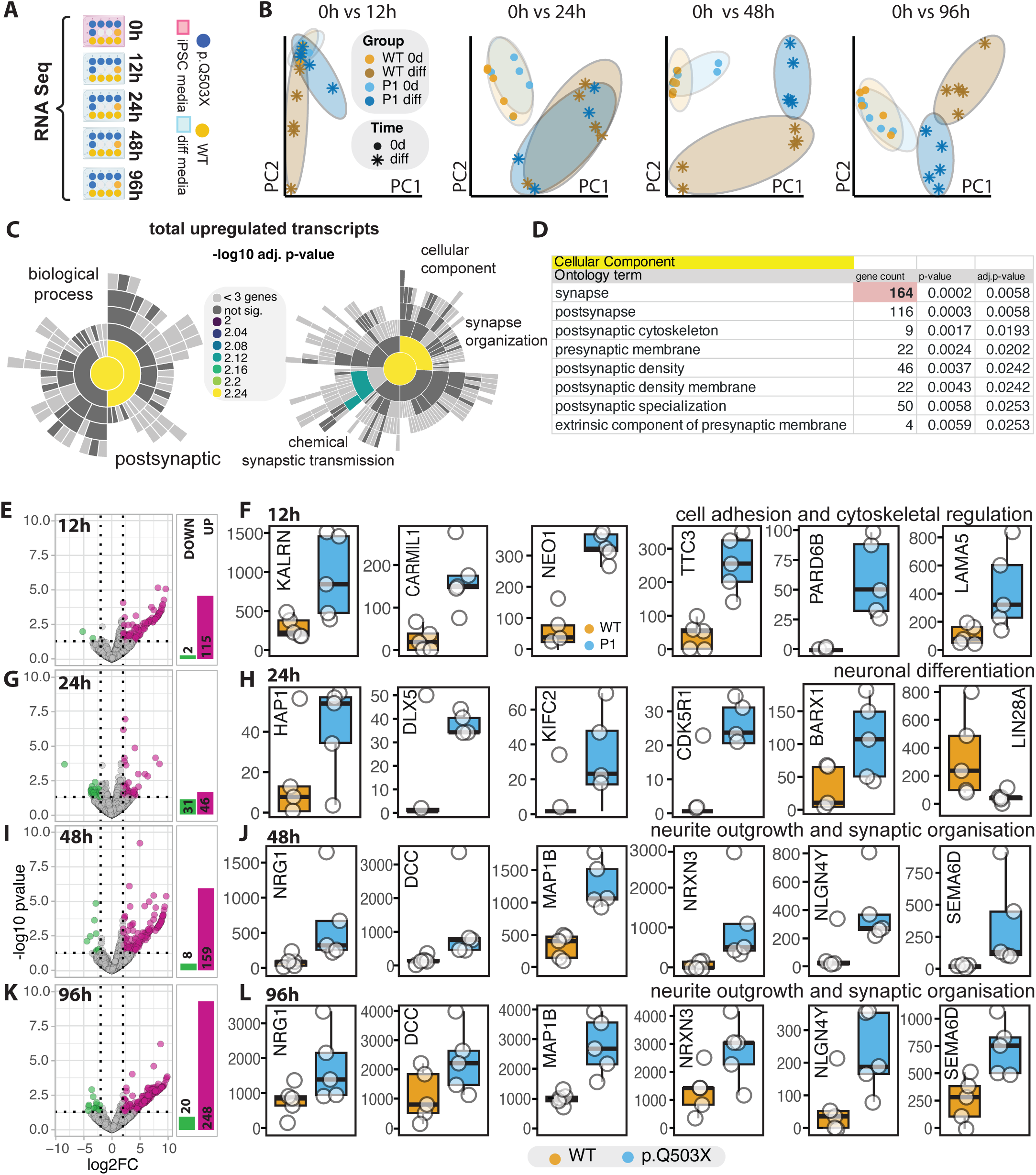
A. Experimental design of RNA-seq analysis. SYNGAP1 p.Q503X haploinsufficient (P1) and isogenic WT iPSCs were collected at 0 h and at 12 h, 24 h, 48 h, and 96 h after induction of GABAergic neuronal differentiation. B. Principal component analysis showing separation between P1 and WT cells across time points. Each panel represents the comparison of 0 h vs 12 h, 24 h, 48 h, or 96 h. C. SynGO enrichment analysis of total upregulated transcripts across all time points. Sun plots show enriched biological process and cellular component categories. Color scale indicates −log10 adjusted p value. D. Table showing representative enriched cellular component categories among upregulated genes, including synapse, postsynaptic membrane, postsynaptic density, postsynaptic cytoskeleton, presynaptic membrane, and postsynaptic specialization, together with gene counts, p values, and adjusted p values. E, G, I, K. Volcano plots showing differential gene expression between P1 vs WT at 12 h (E), 24 h (G), 48 h (I), and 96 h (K). Upregulated genes are shown in magenta and downregulated genes in green (|log2FC| > 2, FDR < 0.05). Numbers of total upregulated and downregulated genes are shown in a barplot. F, H, J, L. Box plots showing representative differentially expressed genes at each time point. F. 12 h: genes related to cell adhesion and cytoskeletal regulation, including KALRN, CARMIL1, NEO1, TTC3, PARD6B, and LAMA5. H. 24 h: pro neuronal differentiation genes including HAP1, DLX5, KIFC2, CDK5R1, ASCL1, and BARX1.J. 48 h: neurite outgrowth and synaptic organization genes including NRG1, DCC, MAP1B, NRXN3, NLGN4Y, and SEMA6D. L. 96 h: neurite outgrowth and synaptic organization genes including NRG1, DCC, MAP1B, NRXN3, NLGN4Y, and SEMA6D.

SynGO analysis of all the upregulated transcripts between the first time points (T=1 to T=4) shows a significant enrichment of transcripts corresponding to synaptic and postsynaptic proteins, in particular those related to proteins that helps to organize synaptic structures and chemical synaptic transmission (Fig. 3C-D). Thus, suggesting an early acceleration of the differentiation to GABAergic neurons.

Across the differentiation time course, the majority of differentially expressed genes in SYNGAP1 haploinsufficient cells were upregulated rather than downregulated, particularly at 12h (115 up, 2 down), 48h (159 up, 8 down), and 96h (248 up, 20 down), while 24 hours showed a more balanced distribution (45 up, 31 down) (Fig. 3E-K). After 12hs post induction, SYNGAP1 haploinsufficient cells showed a dysregulation of cell adhesion, and regulators of cytoskeletal function. In particular, processes associated to the regulation of RHO GAP/GEF/PAK function and lamellipodial protrusions and actin polymerization. This included the upregulation of KALRN, CARMIL1, NEO1, TTC3, PARD6B, and LAMA5 (Fig.3E-F and Suppl. Table 3). Within 24hs post induction, SYNGAP1 haploinsufficient cells shows a dysregulation of pro-neuronal transcripts including, HAP1, DLX5, KIFC2, CDK5R1, and BARX1 (Fig.3F-G and Suppl. Table 3). At this stage, we also determined a significant decrease in LIN28A (Fig.3F-G and Suppl. Table 3), which is kept downregulated at subsequent time points (Suppl. Table 3). LIN28A- mediated let-7 inhibition is required for embryonic development and maintenance of the pluripotent state. Through the inhibition of maturation of let-7 microRNAs, LIN28A inhibits cell differentiation and promote proliferation^35,36^. Within the first 48hs, an increase in neurite outgrowth and synaptic organization transcripts are upregulated in SYNGAP1 mutant cells including NRG1, DCC, MAP1B, NRXN3, NLGN4Y, and SEMA6D (Fig 3I-J). This suggests that within 24-48hs post induction, SYNGAP1 haploinsufficient cells are already set on an accelerated path on neuronal differentiation (Fig. 3I-J; Suppl. Table 3). Importantly, transcripts associated with neurite outgrowth and synapse formation remain upregulated up to 96h, with further increases in absolute gene expression levels compared to 48 hours indicating enhanced activation of neuronal maturation programs (Fig 3K-L).

### Biochemical characterization of SYNGAP1 p.Q503X (patient) GABAergic neurons

We first detected SST positive neurons at 7DIV in SYNGAP1 haploinsufficent neurons, and equal numbers of differentiated SST neurons in p.Q503X (patient) and its correspondent isogenic control. Therefore, to determine the effects of the accelerated neuronal differentiation in SYNGAP1 haploinsufficient neurons, we analyzed total protein levels of whole neuronal extracts for SYNGAP1 p.Q503X (patient) and its correspondent isogenic control SYNGAP1 p.Q503X-c (patient) at 21DIV post neuronal induction. We used four replicate assays per genotype and performed Mass Spectrometry (MS) analysis of iNs as we previously described^33^. We detected changes in protein expression in 1525 proteins with an increase in 776 proteins and a decrease in 689 proteins (sca.adj.pval <0.01) (Fig. 4 A. Suppl. Table 4). Because we observed an increase in mushroom spines (Fig.2), we asked if these morphological changes were also reproduced at the total proteome level. Therefore, we performed SynGO analysis for upregulated and downregulated proteins, using the total GABAergic iNs proteome as a background control. This analysis shows a significant upregulation in protein levels for different components of the presynaptic and postsynaptic signaling machinery, together with synaptic scaffolds such as DLGs, DLGAPs, CASK, MPPs, GEPHYRIN (Fig. 4 B. Suppl. Table 4,5). The increase in these protein levels is reflected on a significant upregulation in structural functions in both, presynaptic and postsynaptic sites (Fig. 4 B. Suppl. Table 5). Moreover, we also determined an increase in protein families corresponding to neurotransmitter release such as Synapsins, Synaptotagmins, SNAPs, or components of synaptic vesicles such as SV2A and VAT1L. These changes occurred together with an increase in total protein levels of proteins involved in GABA metabolism and transport such as ABAT, SLC32A1 (VGAT), neurite growth, synaptogenesis and synaptic transmission (Fig. 4 B. Suppl. Table 4,5). In contrast, downregulated proteins did not include changes in any of these components. Downregulated proteins were mostly enriched in metabolic proteins and DNA/RNA binding, and in proteins regulating transcriptional processes, specifically those involved in neuronal development such as YY1, CIZ1, LIN28A and ASCL1 (Fig. 4 C. Suppl. Table 4). Altogether, this shows that changes in neuronal and synaptic development observed at the morphological level (Fig. 2A-D), are also evidenced at the molecular level, and suggests that a decrease in SYNGAP1 total protein level produces enhancement in the synaptic transmission machinery of human GABAergic neurons.

**Figure 4.**
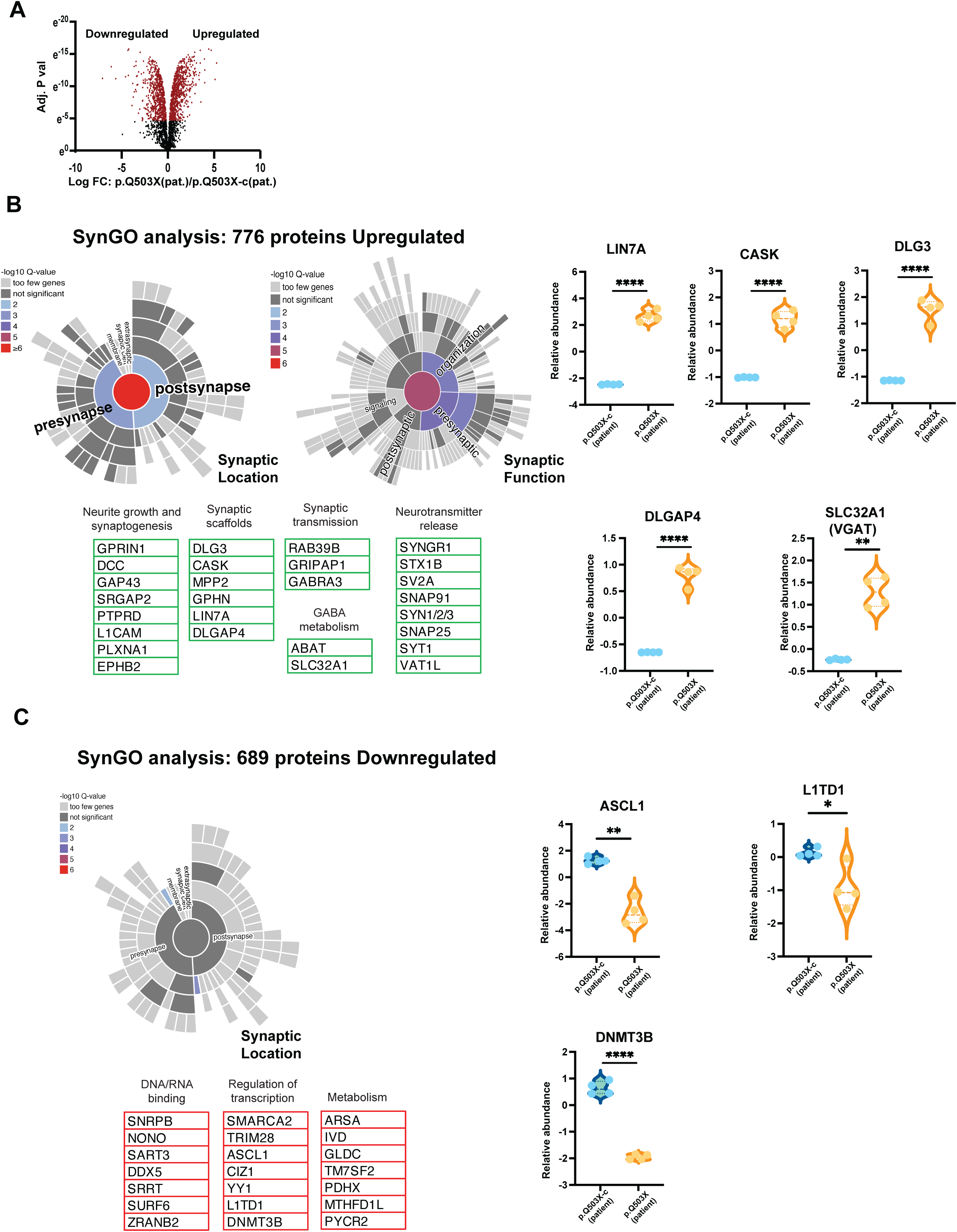
A. Total cell extracts from 21 DIV GABAergic neurons from p.Q503X (patient) and its correspondent isogenic control p.Q503X corrected (P.Q503X-c (patient)) were analyzed by Mass Spectrometry in 4 replicates per genotype (sca.adj.pval <0.01) (A) shows volcano plot with upregulated and downregulated proteins in p.Q503X GABAergic iN. B and C. SynGO analysis of upregulated (B) and downregulated (C) proteins shows a significant increase in 776 synaptic (pre and post) proteins in SYNGAP1 haploinsufficient model. Upregulated proteins are significantly enriched in Neurite growth, synaptogenesis, synaptic scaffolds, synaptic transmission machinery and neurotransmitter release. Components of GABA synthesis and GABA metabolism were also upregulated. No synaptic components were downregulated. Proteins with decreased expression levels were associated to developmental processes, DNA/RNA binding, metabolism and modulators of transcription (lower panel). B and C. Violin plots show quantitative examples of pre/postsynaptic scaffold, GABA transporter VGAT upregulated (B) and downregulation of developmental proteins (C).

### SYNGAP1 PDZ ligand motif is necessary for normal development of human GABAergic neurons

It has been shown that the Syngap1 protein and Syngap1𝞪1 isoform function as a negative regulator for excitatory synapse structure and mechanisms in rodent (cellular and animal) models^17,25^ ^18,37,38^. Moreover, it is reported that a mutation in the PDZ ligand motif of Syngap1 that prevents the binding to PDZ containing proteins is enough to reproduce the impairment in learning phenotypes observed in Syngap1 haploinsufficiency^13^. However, it is not known if the SYNGAP1 PDZ motif is necessary for the normal development of human Glutamatergic or GABAergic neurons. Thus, here we generated a novel iPSC line with a double point mutation in the PDZ ligand motif of the SYNGAP1𝞪1 isoform replacing the c-terminal sequence QQTRV with QQIRE. We have previously shown that this mutation was enough to disrupt the Syngap1 PDZ ligand function, generating a mouse model of Syngap1 dysfunction^13^. Suppl. Figure 1A shows the generation and characterization of the SYNGAP1 PDZ ligand-mutant iPSC line on the WT(03231) background [SYNGAP1 T1306I/V1308E (PDZ Ligand; 03231)] previously used to reproduce the SYNGAP1 p.Q503X haploinsufficient mutation [SYNGAP1 p.Q503X (03231)] (Fig. 2). To determine the correct generation of the desired mutation we sequenced, and karyotyped (Suppl. Figure 1B) the novel cell line and determined the protein levels of SYNGAP1 in GABAergic iNs. For this purpose, we used one of our designed SYNGAP1 specific peptides (^493^AIEEMRLIGQK^504^)^20^ and quantitated SYNGAP1 protein levels in the normal WT(03231) cell line and in the SYNGAP1 PDZ- QIRE(03231) line. Here we used the AIEEYMRLIGQK peptide, to quantitate SYNGAP1 total protein in the WT(03231) and PDZ-QIRE(03231) cell lines by MS. This assay showed that the mutation T1306I/V1308E in SYNGAP1 did not change SYNGAP1 total protein levels (Suppl. Figure 1C-D).

We then asked if the mutation of the PDZ ligand motif can dysregulate neuronal and synaptic maturation of human GABAergic neurons. We first compared the PDZ- QIRE(03231) cell line with its isogenic control, WT(03231), together with the SYNGAP1 p.Q503X haploinsufficient mutation in the same 03231 background [SYNGAP1 p.Q503X(03231)]. Fig. 5 shows that the mutation in SYNGAP1 that impairs its capacity to associate to PDZ domains is enough to reproduce the effects of the SYNGAP1 p.Q503X haploinsufficient mutation. Fig. 5 A-C show that SYNGAP1 PDZ mutant develops neurons with larger dendrites, increases the spine density and shows more spines with a significant increase in mushroom spines. We also observed no significant differences between the SYNGAP1 PDZ-QIRE (03231) and the haploinsufficient SYNGAP1 p.Q503X (03231) genotype.

**Figure 5.**
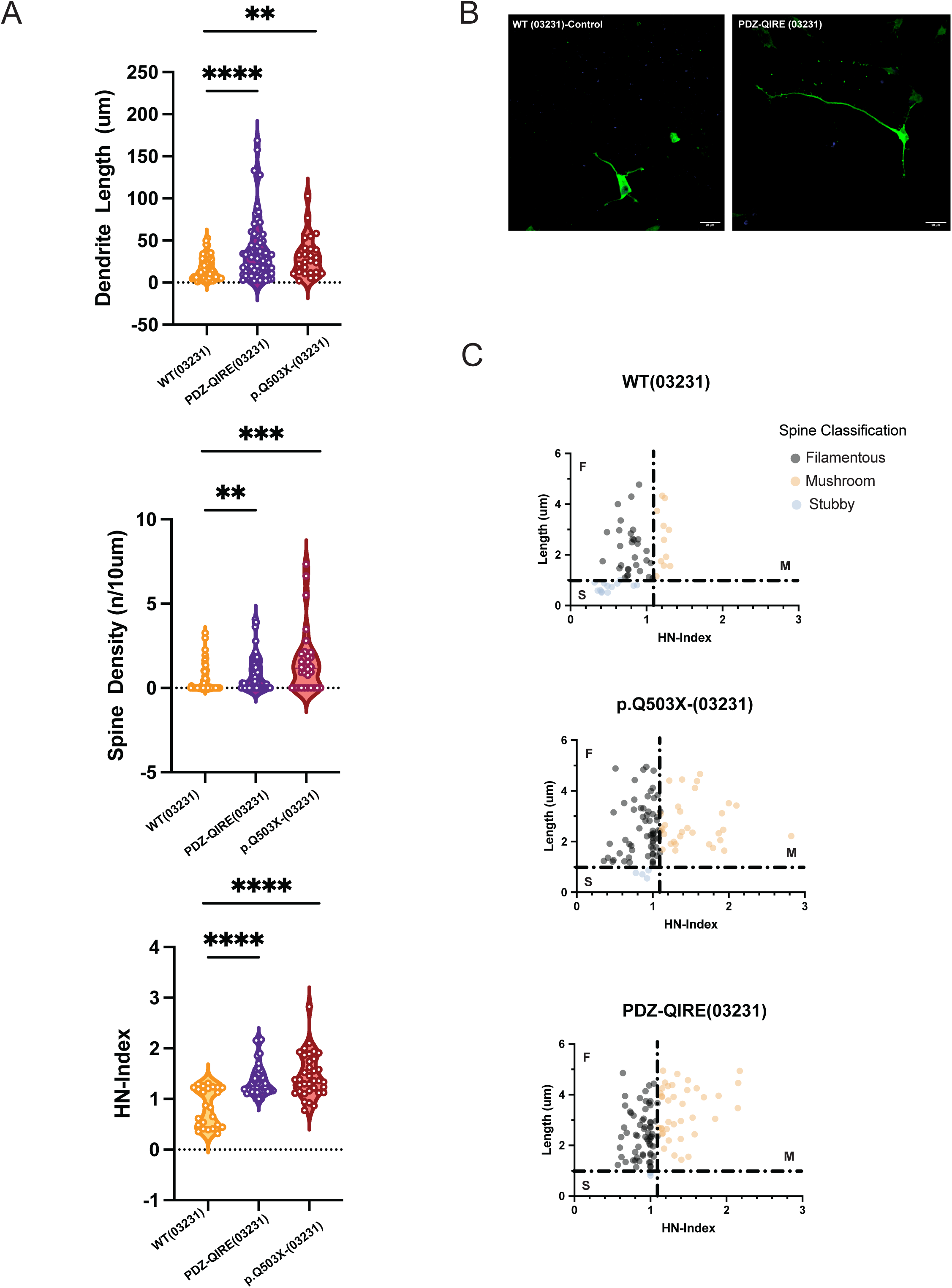
A. Quantification of dendritic length, spine density and HN-Index in WT (03231), PDZ- QIRE (03231) and patient mutation p.Q503X in the 03231 background. The two mutants and control cell line have the same genetic background. Violin plots show that the lack of capacity of SYNGAP1 to associate to PDZ containing proteins in the PDZ-QIRE mutant line generates GABAergic neurons with larger dendrites (p<0.0001), increased spine density (p=0.0042) and more mature spines as shown by a significant higher HN-index (p<0.0001). Similar results are observed in the haploinsufficient SYNGAP1 mutation p.Q503X (03231). The number of spines was determined as spines per 10um of 40-80um dendritic segment. For each cell line, 6 replicates were plated from at least three differentiations. B. Representative confocal max intensity projection images of cultured GABA-iNs at 9 DIV after GFP-transfection at 6DIV from WT (03231) cell line and corrected cell line and PDZ-QIRE mutant SYNGAP1 (scale bar 20um). C. Individual non-filopodial spines plotted in accordance with their length (um) and HN- Index for each line. Figure shows an increase in the number of filamentous and mushroom spines in GABAergic iN, in the SYNGAP1 PDZ mutant cell line and the haploinsufficient p.Q503X (03231), compared to their correspondent isogenic control. Two-sided Mann- Whitney testing was used. Differences between more than two groups were analyzed by one way-ANOVA with Tukey correction for multiple testing, unless the data was non- normally distributed for which a Kruskall-Wallis H test was used. Error bars represent the s.e.m.

We then wanted to determine if the PDZ ligand mutation produced changes in the protein composition of GABAergic synapses. Therefore, we performed a MS analysis of SYNGAP1 PDZ-QIRE(03231) GABAergic neurons and their correspondent isogenic control, WT(03231). Fig. 6 A shows that the SYNGAP1 PDZ-QIRE(03231) mutation produced an increase in the total levels of pre and postsynaptic proteins (Suppl. Table 6). As observed in the SYNGAP1 haploinsufficient model (Suppl. Table 4), downregulated proteins did not include pre and postsynaptic proteins (Fig. 5 B. Suppl Table 6). The removal of SYNGAP1 PDZ ligand increases the levels of synaptic scaffolds including DLG3, DLGAP4, CASK, GPHN, LIN7A; synaptic adhesion molecules as NLGNs/NRXNs; molecules that regulate synaptic growth and synaptogenesis such as ROBO proteins, DCC, GAP43; GABA metabolism: ABA, SLC32A1 (VGAT) and synaptic vesicle and neurotransmitter release such as PCLO, SV2A, SYP, Synapsins, Synaptotagmins, and SNAPs (Fig. 6 A. Supp. Table 6). As observed with a decrease in SYNGAP1 total protein levels, downregulated proteins included enzymes involved in metabolic processes, DNA/RNA binding proteins and transcriptional regulators such as, RCOR2, CIZ1 (Fig. 6 B, Suppl. Table 6). However, within the downregulated proteins there are no major synaptic components. A comparative analysis of dysregulated proteins in the haploinsufficient and PDZ ligand mutant models allowed us to match 992 proteins between datasets, with 416 common proteins being upregulated and 576 downregulated proteins in the SYNGAP1 PDZ-QIRE(03231) mutant genotype (Fig. 6 C. Suppl. Table 6). This comparison shows that 85% of the upregulated proteins are shared within both genotypes, while 97% of the downregulated proteins were also reproduced between the haploinsufficient and the PDZ mutant models (Fig. 6 C. Suppl. Table 6). This shows that the lack of binding to PDZ domains of SYNGAP1 produces a similar protein composition profile to a SYNGAP1 haploinsufficiency model.

**Figure 6.**
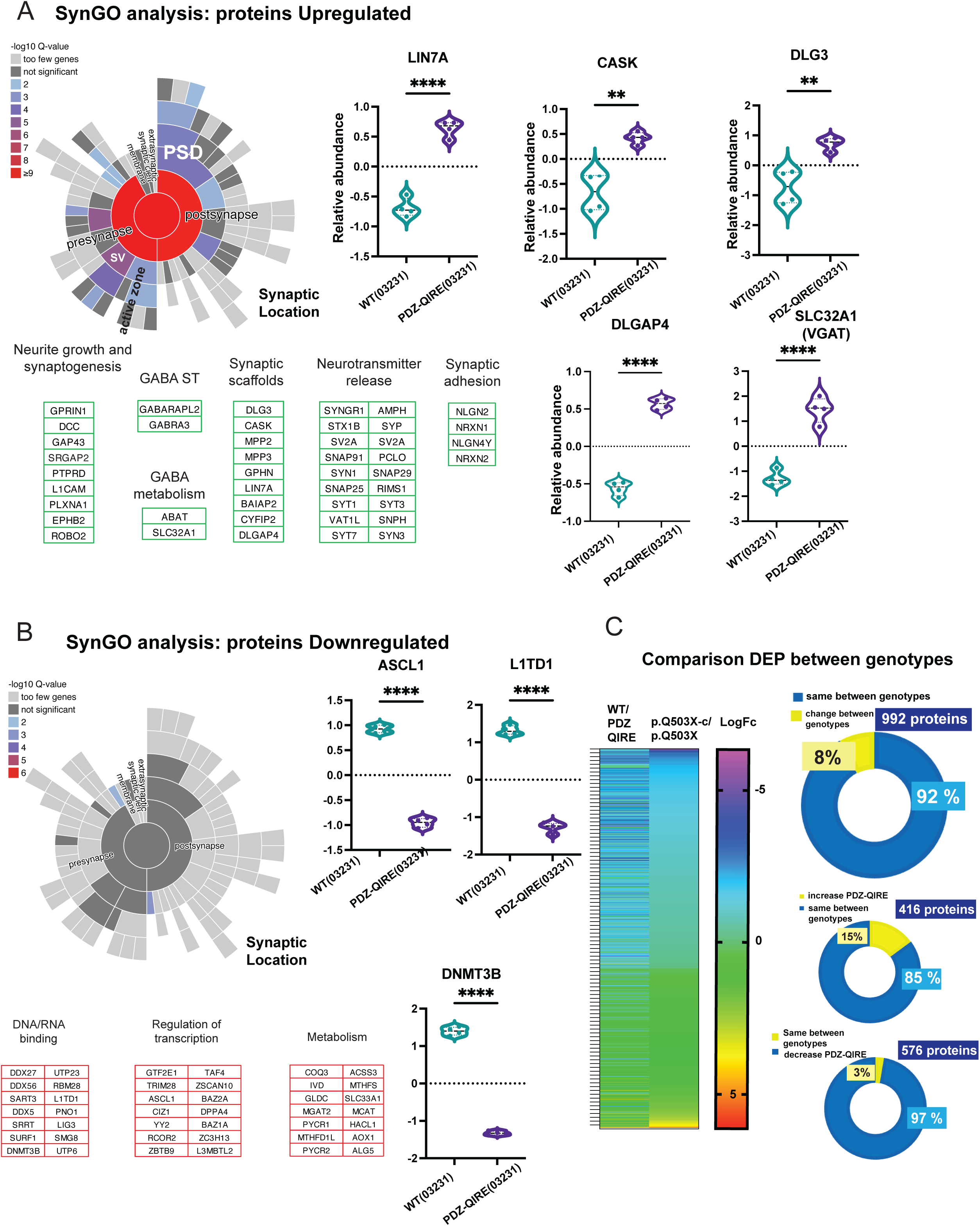
A-B. Total cell extracts from 21 DIV GABAergic neurons from SYNGAP1 PDZ mutant cell line (PDZ-QIRE-(03231)) and its correspondent isogenic control WT (03231) were analyzed by Mass Spectrometry, with 4 replicates per genotype (sca.adj.pval <0.01). A. SynGO analysis (sun plot) shows that upregulated proteins are significantly enriched in components of the pre and post synapse, including postsynaptic density, synaptic vesicles, active zone and structural components of the pre and postsynaptic sites and integral components of their membrane compartments. Violin plots show quantitative examples of pre/postsynaptic scaffolds proteins, and GABA transporter VGAT upregulated in PDZ-QIRE (03231) GABAergic iN. As observed in the p.Q503X SYNGAP1 mutations, upregulated proteins are significantly enriched in Neurite growth, synaptogenesis, synaptic scaffolds, synaptic transmission machinery, synaptic adhesion, and neurotransmitter release. Components of GABA synthesis, metabolism and GABA synaptic transmission were also upregulated. B. Downregulated proteins are depleted of synaptic proteins. Proteins with decreased expression levels were associated to developmental processes, DNA/RNA binding, metabolism and modulators of transcription (lower panel). C. Comparison of upregulated, and downregulated proteins quantitated in the PDZ-QIRE (03231) mutant model and p.Q503X(patient) mutant cell line, compared against their correspondent isogenic controls. From a total of 992 proteins dysregulated in both genotypes, 92% of proteins occurred with the same direction, with 416 proteins upregulated and 576 proteins downregulated, both in the PDZ-QIRE (03231) and the p.Q503X(patient) genotypes. Heat map shows comparison of upregulated/downregulated proteins (Log fold change) for 992 identified proteins between WT (03231)/PDZ-QIRE (03231) and p.Q503-corrected(patient)/p.Q503X(patient). Similar color code indicates same direction in the fold change. Results show that the lack of PDZ-ligand capacity of SYNGAP1 reproduce the molecular changes observed in the haploinsufficient model of SYGAP1 dysfunction.

The increase in total protein levels corresponding to synaptic proteins was also evidenced by increases in protein phosphorylation. We determined protein phosphorylation in the same total extracts from the SYNGAP1 PDZ-QIRE(03231) iNs and their correspondent isogenic control, WT(03231) by MS. We quantitated changes in 1957 phosphopeptides within 1256 proteins, however, a large fraction of these changes corresponded to proteins that also have an increase in total protein levels. This includes 867 dysregulated phosphopeptides matching dysregulated proteins (Suppl. Fig. 2 Suppl. Table 6 and 7). However, a set of 1088 upregulated phosphorylation sites occurred within 797 proteins with no changes in their total protein amount (Suppl. Fig. 2, Suppl. Table 7). Thus, these phosphorylation sites can indicate signaling processes disrupted in the SYNGAP1 PDZ- QIRE iNs because alterations in their signaling components and not because of their change in their protein level. Analysis of these phosphorylation sites shows a dysregulation in several synaptic (p=7.9e-8) processes including the postsynaptic density complex (postsynapse p=1.45e-7, PSD p=3.00e-6) (Suppl. Fig. 3 Suppl. Table 8). This set of 117 synaptic proteins include components of the SYNGAP1 interactome such as ANKS1B, ANK3, CYFIP1, RNA binding proteins and modulators of GTP signaling together with presynaptic proteins. (Suppl. Fig. 3. Suppl. Table 8). Dysregulation of non-synaptic functions was enriched in mechanisms regulating mRNA processing, and transcriptional control (Suppl. Fig. 3 B. Suppl. Table 9). These findings suggest that synaptic signaling, including GTP cascades and SYNGAP1 PPIs, together with processes regulating splicing and RNA post transcriptional control, are the main processes affected by the impairment of SYNGAP1 function in developing GABAergic iNs.

### Regulation of neuronal network activity by impairments in SYNGAP1 function

Our results shows that SYNGAP1 haploinsufficiency or the lack of SYNGAP1 to associate to PDZ containing proteins, produces an acceleration of neuronal maturation that generates neurons, with an increase in the number of more mature spines. Therefore, to determine if these biochemical and morphological changes can produce changes in neuronal activity, we generated co-cultures of Glutamatergic and GABAergic iNs (Fig. A). We used a ratio 80/20 Glutamatergic/GABAergic, since this ratio have been proposed to better represent the neuronal composition of developing cortical circuits in different in vitro models and in vivo^39,40–42^. It has been reported that GABAergic iNs became functionally inhibitory after 42DIV^43^. We then confirmed the inhibitory GABAergic signaling expressed in co-cultures, by comparing the mean spike frequency of glutamatergic neurons cultures with and without GABAergic neurons at DIV 56. In these conditions, we observed a significant inhibitory control of GABAergic iN (Fig. 7B). As a control, we used a co-culture of WT (03231) Glutamatergic neurons with the same genotype WT (03231) used for GABAergic neurons. We then compared the mean spike frequency of cultures using the p.Q503X-c (patient corrected) genotype glutamatergic neurons co-cultured with the same genotype (p.Q503X-c) GABAergic neurons, against co-cultures with p.Q503 X (patient mutation) GABAergic neurons (Fig. 7B). We determined that co-cultures using GABAergic neurons carrying the haploinsufficient SYNGAP1 mutation p.Q503X, produced a significantly larger decrease of the mean spike frequency, when compared against co-cultures with its isogenic control (Fig, 7B). We then performed the same assay using the SYNGAP1 PDZ ligand mutant cell line PDZ- QIRE(03231) and compared co-cultures against their isogenic control WT (03231). In both cultures, excitatory neurons were generated from the isogenic WT (03231) cell line (Fig. 7B). With this setup, we observed a similar result as shown for SYNGAP1 p.Q503X, with a significant decrease in the mean spike frequency of co-cultures when compared to its isogenic control (Fig. 7B). Therefore, these experiments show that a decrease in SYNGAP1 total protein levels or a lack of PDZ ligand function is enough to disrupt neuronal network activity in mixed Glutamatergic-GABAergic co-cultures. All recordings were performed using 8 by 8 grids multielectrode arrays (MEAs), after 56DIV (Fig. 7A).

**Figure 7.**
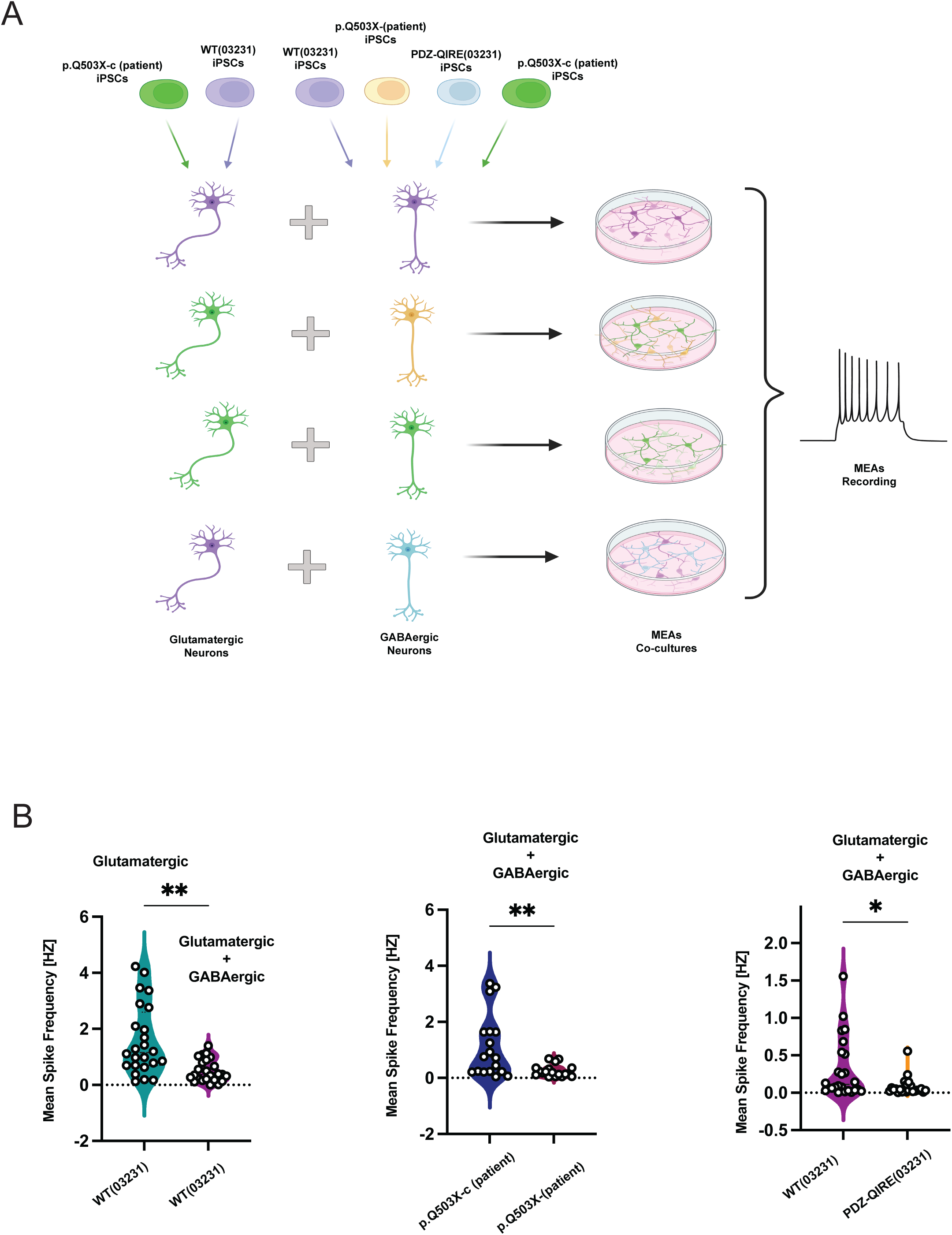
A. Diagram of the experimental pipeline, Glutamatergic iN where co-cultured with GABAergic neurons in a ratio 80/20 for 8 weeks and the mean spike frequency recorded using a MultiChannel Systems MEA-2100 multielectrode array (MEA). Glutamatergic neurons induced via NGN2 expression from WT (03231) iPSC cell line or patient corrected p.Q503X-c (patient) line, where used in cultures in combination with WT (03231) GABAergic neurons, patient derived p.Q503X (patient) GABAergic neurons, patient corrected p.Q503X-c(patient) GABAergic neurons, or PDZ-QIRE (03231) GABAergic neurons. All the cell lines were compared against their correspondent isogenic controls. B. Left violin plot shows a significant decrease in the Mean spike frequency of the recordings when WT (03231) glutamatergic neurons were co-cultured with WT (03231) GABAergic iN neurons in comparison to glutamatergic WT (03231) monocultures P=0.0228 [Welch’s t test]. Middle plot shows recordings from co-cultures of patient corrected p.Q503X-c (patient) glutamatergic neurons with the same patient corrected p.Q503X-c (patient) GABAergic neurons (control recording) compared to p.Q503X-c (patient) glutamatergic neurons co-cultured with the patient mutation p.Q503X (patient) GABAergic iN. Figure shows a significant decrease in the mean spike frequency in co-cultures corresponding to the p.Q503X haploinsufficient mutation P=0.0259 [Kolmogorov-Smirnov test]. Right violin plot shows that similar results P=0.0307, [Welch’s t test] are obtained when the control cell line WT (03231) co-cultures (glutamatergic + GABAergic) are compared to WT (03231) Glutamatergic neurons co-cultured with PDZ-QIRE (03231) GABAergic neurons, suggesting that both the p.Q503X haploinsufficient mutation and PDZ mutant cell line produces similar effects on the recorded network activity. Results corresponds to 6 replicates and three independent differentiations.

## Discussion

Different lines of evidence suggest a role of SYNGAP1 at early stages of human neuronal development^20,21^. However, the role of different SYNGAP1 domains and the function in different neuronal types, particularly the role of SYNGAP1 isoforms in development, are not clear. Here we show that SYNGAP1 is not a specific decelerator of the differentiation of Glutamatergic neurons but has a broader effect that also includes GABAergic neurons. Here, we show that SYNGAP1 role in neuronal differentiation starts at the very early stages of neuronal induction with a clear transcriptional dysregulation setting SYNGAP1 mutant cells into a neuronal path within the first 12hs post neuronal induction. Transcriptional analysis suggests an early dysregulation of cell adhesion and cytoskeletal processes that follows with transcriptional processes controlling neuronal differentiation. We determined an early downregulation of LIN28A transcripts as early as 24hs post neuronal induction, that were also confirmed at neuronal stages by MS analysis. LIN28A promotes proliferation and prevents neuronal differentiation^36^. This pro-neuronal is followed with an upregulation of axon guidance, synaptic cell adhesion and scaffold transcripts in SYNGAP1 mutant cells. This accelerated differentiation in then followed with a significant increase in the identification of SST positive neurons at 7DIV. At this developmental stage, no electrical activity is detectable by MEAs. By 21DIV, haploinsufficient neurons have larger dendrites, more spines and a larger fraction of mushroom spines. MS assays shows that these spines contains a significant increase in synaptic proteins, as well as in components of GABA transport, synthesis and its signaling machinery. The increase in these proteins co-occurred with a significant downregulation in proteins associated to immature and proliferative processes, including LIN28A. At 56DIV, these show a stronger GABAergic inhibitory tone in co-cultures of WT Glutamatergic and mutant GABAergic neurons, evidenced by a reduction in their spike frequency. This is likely because of an increase in the activity of the more mature GABAergic neurons. Our results shows that an intact PDZ ligand motif is essential for SYNGAP1 to function as a decelerator of neuronal differentiation. This suggests that the precise localization of SYNGAP1𝞪1 to its interacting partners is critical for its function at very early stages of neuronal maturation. Thus, the capacity of SYNGAP1 to associate to proteins containing PDZ domains seems to be essential for its role in controlling the speed of differentiation of GABAergic neurons. In human radial glia cells, SYNGAP1 co- localizes at tight junctions with the PDZ and MAGUK family member TJP1^20^, suggesting that a PDZ mediated protein interaction might also be necessary. While comparing between different models and conditions is difficult, it seems that the capacity of SYNGAP1 to associate to proteins containing PDZ domains is a necessary condition for normal SYNGAP1 function as a regulator of neuronal development and synapse structure. We expect that a decrease of SYNGAP1 protein levels or the lack of PDZ binding capacity will impair synaptic function. However, here we focus on the role of SYNGAP1 and its PDZ ligand function at early neuronal differentiation and not in its synaptic function. We show that SYNGAP1 PDZ ligand function is essential as a decelerator of neuronal (GABAergic) differentiation independently of its synaptic role. Future studies will address if SYNGAP1 mutation in its PDZ ligand domain impairs its capacity to associate to TJP1 and other PDZ containing proteins at hRGCs tight junctions, and what are the protein interaction networks that are dysregulated by the impairment of SYNGAP1 association to PDZ containing proteins.

## Methods

### Cell Lines, Cell Culture and Neural Differentiation

The iPSC lines were generated using Yamanaka factors in patient-derived PBMCs, as previously described (17). iPSC lines were maintained with mTeSR plus (STEMCELL Tecnologies #100-0275) media changes every other day on 1:200 geltrex (GIBCO, #A1413301) coated tissue culture 6-well plates (Genesee Scientific #25-105) and passaged using ReLeSR (STEMCELL Technologies #05872). Cells were maintained below passage 50 and periodically karyotyped via the G-banding Karyotype Service at Children’s Hospital Los Angeles.

### CRISPR/Cas9-edited iPSC Line Generation

#### SYNGAP1 p.Q503X Corrected Line

CRISPR/Cas9 editing technology was utilized to correct the patient nonsense (c.1507C>T; p.Q503) mutation and generate an isogenic corrected control line p.Q503X-c (patient). A sgRNA targeting the patient-specific mutation (p.Q503X) in *SYNGAP1* was cloned into pSpCas9(BB)-2A-Puro (PX459) V2.0 (Addgene plasmid #62988). The sgRNA construct and the HDR template containing the WT *SYNGAP1* sequence, were electroporated into the p.Q503X patient iPSC line. Individual iPSC colonies were transferred to 24 well plates and subsequently underwent restriction enzyme-based genotyping. Positive colonies were then confirmed via Sanger sequencing and expanded in culture.

HDR Template: CCGCGAGAACACGCTTGCCACTAAAGCCATAGAAGAGTATATGAGACTGATTGGT<u>C</u> <u>AG</u>AAATA<u>T</u>CTCAAGGATGCCATTGGTATGGCCCACACTCAGGCCCTCTTCTTCCCAA ACCTGCCA

The underlined CAG sequence corresponds to the c.1507 insertion of the WT “C” base pair. The substitution of the truncating “T” with the WT “C” base pair was screened for via restriction enzyme digestion (DrdI) and then confirmed via sanger sequencing. The underlined T base corresponds to a silent PAM mutation as we previously reported^20^.

#### *SYNGAP1* Mutation Insertions

The SYNGAP1 p.Q503X (03231) cell line was generated via substitution of c.1507C>T; p.Q503X. A sgRNA targeting the *SYNGAP1* c.1507C site was cloned into pSpCas9(BB)-2A-Puro (PX459) V2.0. The sgRNA and the HDR template were nucleofected in the WT (03231) control line. Individual iPSC colonies were transferred to 24 well plates and subsequently underwent restriction enzyme-based genotyping. Positive colonies were then confirmed via sanger sequencing.

HDR template: CCGCGAGAACACGCTTGCCACTAAAGCCATAGAAGAGTATATGAGACTGATTGGT<u>TA</u> <u>G</u>AAATA<u>T</u>CTCAAGGATGCCATTGGTATGGCCCACACTCAGGCCCTCTTCTTCCCAAA CCTGCCA

The underlined TAG region shows the c.1507C>T inserted mutation and the underlined T base shows the introduced silent PAM mutation. The substitution of the WT “C” with the truncating “T” base pair was screened for via restriction enzyme digestion (DrdI) and then confirmed via sanger sequencing as previously reported^20^.

The SYNGAP1 PDZ QRTV-QIRE mutation was generated, characterized and the insertion confirmed by Synthego-EditCo Bio, Redwood City, CA, USA.

### Neural Differentiation

#### GABAergic Induced Neurons

iPSC cells were seeded at a density of 1.5×10^5^ cells/well on geltrex-coated 6-well plates with mTesR Plus media and 10 μM Rock inhibitor Y-27632. The following day, cells were infected with lentivirus plasmid with an inducible expression of ASC11 and constitutively expresses the Puromycin resistance gene (packaged using pTet-O-FUW-Asc11- puromycin plasmid from Addgene, Plasmid #97329 (Addgene, Cambridge, MA)), lentivirus plasmid that inducible expresses DLX2 and constitutively expresses the Hygromycin resistance gene (packaged using pTet-O-FUW-Dlx2-hygromycin plasmid from Addgene, Plasmid #97330 (Addgene, Cambridge, MA)), and a reverse tetracycline transactivator, 1 μg/ml of doxycycline and 2 μg/ml of polybrene. After overnight incubation, the media was replaced with fresh mTeSR Plus media with 2 μg/ml of Doxycycline. The next day, 0.7 μg/ml of puromycin was added to the culture. After a 48hr selection, the media was changed with fresh mTeSR Plus media and cells were fed every other day until they reached approximately 80% confluency. To start neuron induction, confluent cells were dissociated with Accutase and plated onto geltrex-coated 6-well plates in GABA-N2 media (DMEMF12, 1X N2 Supplement, 1X NEAA, supplemented with 10uM of ROCK inhibitor and 1ug/ml doxycycline at a cell density of 2×10^5 cells per well (Day 0). On Day 1, media was replaced with GABA-N2 media supplemented with 1ug/ml doxycycline, 0.7ug/ml puromycin, and 100ug/ml hygromycin (Corning #30-240-CR). On Day 3, media was replaced with GABA-B27 media (Neurobasal, 1X B27 supplement, 1X GlutaMAX) supplemented with 1ug/ml doxycycline and 2uM Ara-C. Media was replaced every other day until cells reached their endpoint.

### Biochemistry

#### Immunoprecipitation

iPSC-derived GABAergic neurons cultured in geltrex coated 10 cm2 tissue culture dishes were harvested via accutase treatment and lysed in cell lysis buffer (50 mM Tris pH 7.4, 2 mM EDTA, 10 mM NaVO4, 30 mM NaF, 20 mM β-glycerolphosphate, 1% n-Dodecyl-β- Maltopyranoside, supplemented with complete Protease Inhibitor Cocktail Tablets using mechanical homogenization followed by incubation at 4 degrees Celsius while rotating for 40 minutes. The cell lysate was then centrifuged at 35,000 RPM for 30 min at 4 degrees Celsius and the protein concentration of the supernatant was subsequently determined using the BCA assay kit (Thermo Cat# 23227). 2 mg of total protein were incubated with anti-SYNGAP1 antibody at 4 degrees Celsius overnight with gentle rotation. The following day, samples were mixed with 50 ul of Dynabeads protein G (ThermoFisher Cat# 10004D) and incubated at 4 degrees Celsius for 2 hours with gentle rotation. Beads were washed 3 times in IP wash buffer (25 mM Tris pH 7.4, 150 mM NaCl, 1 mM EDTA, 1% n-Dodecyl-β-Maltopyranoside) and then incubated with 50 ul 2X LDS sample buffer (Invitrogen Nupage NP0008) on a thermomixer (Eppendorf Cat #) for 15 min at 95 degrees Celsius with rotation (500 rpm). If the samples were for Mass Spectrometry, they were incubated with 10mM DTT (Invitrogen. Cat#15508-0) at 56 degrees Celsius for 1 hour followed by 20mM Iodoacetamide (Sigma I1149-25G) at room temperature for 45 min in the dark.

#### Western Blot

Samples were combined with LDS sample buffer 10 mM DTT and then incubated at 95 degrees Celsius for 15 minutes. Samples were then loaded on 4 – 12% Bis-Tris gels (NuPAGE Novex, Thermo Fisher Scientific, Waltham, MA) and separated at 135V for 1.5 hours. Proteins were then transferred to a PVDF membrane using a Bio-Rad Trans-Blot Turbo Transfer System (Bio-Rad, Hercules, CA). Membranes were blocked for 1 hour at room temperature with 5% bovine serum albumin (BSA) in 0.05% TBS-Tween (TBST) and then incubated with primary antibody overnight at 4 degrees Celsius. Membranes were washed with 0.05% TBST four times, ten minutes each, and then incubated with secondary antibodies for 1 hour at room temperature. Membranes were washed with 0.05% TBST 4 times, 5 minutes each, and imaged using a 4000MM Pro Image Station (Carestream, Rochester, NY).

#### Liquid Chromatography Mass Spectrometry Sample Preparation

SYNGAP1 p.Q503X, p.Q503X-c PDZ-QIRE (03231) and WT (03231) iPSCs were differentiated to GABAergic neurons for two weeks. Pellets were collected and flash-frozen in liquid nitrogen. Samples were processed in at least four replicates from two independent differentiations per each genotype, for total proteome and phosphoproteome analysis or previous PSD preparations before MS assays.

Total neuronal extracts PSD fractions and phosphopeptide enrichement were prepared following the same protocol as described before^33^. Briefly iPSC derived neurons were cultured for 3 weeks we isolated PSDs or total fractions from four replicates per genotype and, performed enrichment of phosphopeptides in total neuronal extracts with titanium dioxide (TiO_2_) chromatography, samples were analyzed using LC-MS/MS. Samples reconstituted in LC buffer A (0.1% formic acid in water), randomized, and then injected onto an EASY-nLC 1200 ultra-high-performance liquid chromatography coupled to a Q Exactive Plus quadrupole-Orbitrap mass spectrometer (Thermo Fisher Scientific). Peptides were separated by a reverse phase analytical column (PepMap RSLC C18, 2 µm, 100 Å, 75 µm×25 cm). Flow rate was set to 300 nL/min at a gradient from 3% LC buffer B (0.1% formic acid, 80% acetonitrile) to 38% LC buffer B in 110 min, followed by a 10-min washing step to 85% LC buffer B. The maximum pressure was set to 1,180 bar, and column temperature was maintained at 50°C. Peptides separated by the column were ionized at 2.4 kV in positive ion mode. MS1 survey scans were acquired at the resolution of 70,000 from 350 to 1,800 m/z, with a maximum injection time of 100 ms and AGC target of 1e6. MS/MS fragmentation of the 14 most abundant ions were analyzed at a resolution of 17,500, AGC target 5e4, maximum injection time 65 ms, and normalized collision energy of 26. Dynamic exclusion was set to 30 sec, and ions with charge +1, +7 and >+7 were excluded. MS/MS fragmentation spectra were searched with Proteome Discoverer SEQUEST (version 2.2, Thermo Scientific) against in silico tryptic digested Uniprot all-reviewed Homo sapiens database. The maximum missed cleavages were set to two. Dynamic modifications were set to phosphorylation on serine, threonine, or tyrosine (+79.966 Da), oxidation on methionine (+15.995 Da), and acetylation on protein N-terminus (+42.011 Da). Carbamidomethylation on cysteine (+57.021 Da) was set as a fixed modification. The maximum parental mass error was set to 10 ppm, and the MS/MS mass tolerance was set to 0.02 Da. The false discovery threshold was set strictly to 0.01 using the Percolator Node. Individual phospho-site localization probabilities were determined by the ptmRS node, and phospho-sites with <0.75 localization probability were removed. The relative abundance of phospho-peptides was calculated by integration of the area under the curve of the MS1 peaks using the Minora LFQ node in Proteome Discoverer. No data imputation was performed for missing values. Phospho- peptides were filtered so that each condition had at least two quantified values. Phospho- peptide intensities were then normalized by log_2_-transformation and sample median subtraction. Only phosphorylation sites with proteins with no changes in the same direction in total protein levels were used for further analysis. For the MS analysis of PSD fractions duplicate samples of the genotype WT 03231 PSD samples were dissolved in 100 µL lysis buffer (0.5M triethylammonium bicarbonate, 1% sodium deoxycholate). Samples were subjected to tip sonication (Q700, QSonica, amplitude = 10, 2 sec on/2 sec off pulses, 20 sec total processing time per sample, on ice) and then centrifuged at 15K rpm at 4^0^C for 10 min. The supernatant was transferred to a fresh tube (on ice) and the protein concentration of each sample was measured using the Qubit Protein Assay Kit (Thermo, Q33211) and Qubit 4.0 fluorometer per manufacturer’s instructions. Equal amount of protein (40 µg) per sample was transferred to a fresh tube adjusted to highest volume (90 µL) with lysis buffer. Four µL Reducing Reagent (Sigma, 4381664) were added to each sample. Samples were incubated at 60^0^C for 1hr. Two µL Alkylating Reagent (Sigma, 4381664) were added to each sample. Samples were incubated at room temperature for 15 min. Two µg trypsin/LysC (Promega, V50703) were added to each sample. Samples were incubated overnight at room temperature in dark reconstituted in water 0.1% formic acid and analyzed using LC-MS with FAIMS ion mobility pre-separation (nano-easy LC 1200, Thermo Orbitrap Exploris 480).

Raw data files of the native peptide LC-MS analysis were submitted to Proteome Discoverer 2.5 (Thermo) for target decoy search using Sequest against the homo sapiens canonical swissprot database (TaxID= 9606, v2021-10-30). The search allowed for up to two missed cleavages, a precursor mass tolerance of 20ppm, a minimum peptide length of six and a maximum of three equal dynamic modifications of oxidation (M), deamidation (N, Q) or phosphorylation (S, T, Y). Methylthio (C) and TMTpro (K, peptide N-terminus) were set as static modifications. Peptide level confidence was set at q<0.05 (<5% FDR).

#### Targeted Parallel Reaction Monitoring-Mass Spectrometry

We used a similar protocol as we previously reported^20^. Briefly, tryptic peptides shared between all SYNGAP1 isoforms as well as peptides unique to the canonical protein and each of the truncated isoforms were synthesized commercially (ThermoFisher Scientific). Both unlabeled light and isotope labeled heavy forms were synthesized. Commercially synthesized labeled peptides used ^13^C and ^15^N labeled lysine or arginine. An internal standard was prepared from the isotope-labeled standards in 3% acetonitrile with 0.1% formic acid with 5 fmol/µL of Peptide Retention Time Calibration (PRTC) mixture (ThermoFisher Scientific) and 10 µg/mL E. coli lysate digest (Waters) as a carrier. Isotope-labeled peptides were dissolved at 2000 pg/mL. Standard Concentration: 1000pg/mL. The selected peptides ^493^AIEEMRLIGQK^504^ correspond to SYNGAP1 (ENSG00000197283) GRCh38.p13,and can be used to quantitate total SYNGAP1 protein levels. Calibration curves were prepared from the light standards, and samples were analyzed by tPRM mass spectrometry using LC-MS with FAIMS ion mobility pre-separation (nano-easy LC 1200, Thermo Orbitrap Exploris 480). The calibration curve used the ratio of light to heavy peptide using a bilinear curve fitting and 1/x^2^ weighting. The limit of quantitation was estimated as the lowest calibration point with a coefficient of variability below 15% and an average error below 15%.

#### Immunofluorescence

Cells were fixed with 4% PFA and permeabilized with 0.1% PBS (Corning #21-040-CV)+ TritonX100 (Sigma #T8532-100ML; PBST) for 15 minutes at room temperature, and subsequently blocked in 1% bovine serum albumin (Genesee Scientific #25-529; BSA) in 0.025% PBST for 2 hours at room temperature and then incubated overnight with primary antibodies in blocking solution at 4C. The following day, cells were washed three times with 0.025%PBST and then incubated with secondary antibodies in blocking solution for 1 hour at room temperature. Cells were triple washed with 0.025% PBST and nuclei were stained with DAPI (1:2000; Cell Signaling #4083) diluted in 1xPBS for 5 minutes. Cells were washed three times for 5 min with 1xPBS before mounting coverslips on slides (VWR #48311-703) with ProLong Diamond Antifade mountant (Invitrogen #P36965).

### RNA sequencing and analysis

RNA seq was performed on SYNGAP1 haploinsufficient p.Q503X lines and isogenic controls across five time points. At T0 iPSCs were collected after reaching confluency in StemFlex media. After cells were seeded in GABA N2 differentiation media, four time points were collected at 12hs (T1), 24hs (T2), 48hs (T3), and 96hs (T4) following with the GABAergic induction and differentiation protocol. RNA was extracted, reverse transcribed into cDNA. Each sample was barcoded using Biostate-AI’s proprietary BIRT technology (Barcode-Integrated Reverse Transcription) + PERD RNAseq technology, and sequenced with the Illumina NovaSeq X Plus platform. Count matrices were analyzed in RStudio using edgeR^44^, similar to previous studies^45–48^. Low expressed genes were removed using filter ByExpr, and libraries were normalized using the TMM method. Differential expression between SYNGAP1 p.Q503X haploinsufficient lines and isogenic controls was performed at each time point using edgeR’s negative binomial exact Test with WT as the reference. Genes with an absolute log2 fold change > 2 and FDR < 0.05 were considered differentially expressed. Principal component analysis was performed on log transformed counts per million values to assess global transcriptional differences across genotypes and time points. Gene set enrichment analysis was conducted using clusterProfiler^6^ gseGO on ranked gene lists based on log2 fold change (with Gene Ontology biological processes). Selected marker genes related to neuronal identity and differentiation were visualized using raw counts across time points and genotypes.

### Functional Assays

#### Dendritic Spine Analysis

Excitatory and inhibitory neurons were cultured as monocultures on poly-L-ornithine (Sigma #P3655-10MG) and laminin (Gibco #23017-015) coated glass coverslips (neuVitro #GG-12-1.5-PRE) in 24 well plates (Falcon #353047) and assayed at 21 and 9 DIV, respectively. For morphological analysis neurons were transfected with GFP (pSin- Ef1a-EGFP Addgene #21320) and spines co-visualized with anti-MAP2 and anti-GFP antibodies. For each assay, morphological analyses of dendritic spines were performed by 3D reconstruction using Imaris Filament Tracer (Bitplane) on images acquired on a Nikon A1R HD microscope with a Nikon Plan Apo 𝛌 60x oil objective. Dendritic length was obtained by manually tracing cells in FIJI on brightfield images acquired on a Nikon DS- Fi3 camera with a Nikon Plan Apo 𝛌 4x objective to capture the entire dendritic field of each cell. All assays were performed in three independent differentiations using 3 replicate assays per differentiation. Data was gathered by an individual blinded to the experimental conditions.

#### Multielectrode array recordings

iPSC-derived glutamatergic neurons were co-cultured with primary murine astrocytes and plated onto six-well multielectrode chips (nine electrodes and one ground per well) coated with poly-L-ornithine and laminin with a ratio 80/20 (glutamatergic/GABAergic). To study the inhibitory effect of GABAergic neurons derived from the different iPSC lines, WT (03231) glutamatergic iNs were co cultured with GABAergic iNs on MEAs. Spontaneous network activity was recorded from day 56 in culture using a MultiChannel Systems MEA- 2100 multielectrode array (MEA) amplifier (ALA Scientific) with built-in heating elements set to 37 °C. Cells were allowed to acclimate for 5 min after chips were placed into the MEA amplifier before beginning 7min recordings. Active electrodes were defined as those that averaged more than 10 spikes per minute. Bursting electrodes were defined as those that had at least 1 burst per minute. Bursting wells were defined as well that had more than 30% bursting electrodes. All wells that did not meet these criteria were discarded. Spikes were detected by crossing of a threshold set to a level of 3 standard deviations from the baseline noise level. Mean spike frequency (Hz), bursts per minute, mean burst duration, and percent spikes in bursts were determined using the accompanying MC Rack software.

#### Data Analysis

Analysis was performed with the statistical software package Prism Origin (GraphPad Software). Differences between two groups were analyzed using a two-tailed Student’s *t* test, unless the data was non-normally distributed for which two-sided Mann-Whitney testing was used. Differences between more than two groups were analyzed by one way-ANOVA with Tukey correction for multiple testing, unless the data was non-normally distributed for which a Kruskall-Wallis H test was used. Significance was assumed at *P* < 0.05. Error bars represent the s.e.m. unless otherwise stated. For MS data analysis, see MS methods.

## Figure Legends

Supplementary Figure 1

A. Generation of SYNGAP1 PDZ ligand mutant cell line. (A) Scheme of SYNGAP1 modular structure with PH, C2, RASGAP, DUF3498 (domain of unidentified function 3498) protein domains, together with the PDZ ligand at the c-terminal region QTRV. The cartoon shows the replacement of c-terminal sequence QTRV, by QIRE.

B. Strategy of the two amino acids substitution and sequence confirmation for the double point mutation of SYNGAP1 PDZ region on the WT (03231) genetic background. iPSC generation showing pluripotent stem cells markers OCT4 and SSE4.

C. Quantitation of SYNGAP1 total levels by MS in NPCs shows no changes in SYNGAP1 total protein levels in SYNGAP1 PDZ mutant compared to its isogenic control, using the SYNGAP1 peptide ^493^AIEEMRLIGQK^504^ for targeted MS analysis.

Supplementary Figure 2

Figure shows SynGO analysis (left) of total number of phosphopeptides upregulated in PDZ-QIRE (03231) GABAergic iN compared to its isogenic control WT (03231). Sun plot shows an enrichment of presynaptic and postsynaptic proteins corresponding to structural, cytoskeletal, membrane components of the PSD together with presynaptic membrane proteins, synaptic vesicle proteins and components of the active zone. Right plots show analysis of synaptic functions enriched within upregulated proteins including organization of the pre and post signaling machinery, assembly of the synapse and synaptic vesicle release. Lower plots show upregulated phosphorylation sites that corresponds only to upregulated proteins. The analysis shows that a large proportion of the observed changes in protein phosphorylation match upregulation in protein levels.

Supplementary Figure 3

A. Analysis of changes in protein phosphorylation curated against changes in protein levels. A total of 1088 phosphopeptides occurred in 797 proteins that were not found to be upregulated in the SYNGAP1 PDZ-QIRE (03231) genotype. A total of 117 of these proteins were identified as synaptic proteins by SynGO analysis. These proteins were enriched in synaptic and postsynaptic components (Sun plot/SynGO), but not presynaptic proteins. Colored circles show different functional groups within the synapse, including components of the SYNGAP1 interactome such as ANKS1B, ANK3, MAPK1, CYFIP1, together with GTP signaling proteins (green), structural and signaling components of the PSD (yellow), presynaptic proteins (blue), RNA binding/ribosomal proteins (purple) protein kinases (pink) and channels (dark blue).

B. Protein phosphorylation sites upregulated in non-synaptic components were enriched in RNA processing, Nucleotide excision repair development, gene expression, cellular growth, and RNA PTMs.

## Acknowledgements

This research was supported by the Syngap Research Fund and NIH R01MH115005 to MPC.

