## Supplementary material for "Disruption of the SYNGAP1 PDZ ligand motif accelerates differentiation of human iPSC-derived GABAergic neurons": Supp Fig 1

### Suppl. Fig1

A

**PDZ ligand mutation:**
SYNGAP1 T1306I / V1308E (PDZ Ligand; 03231) or PDZ-QIRE (03231)

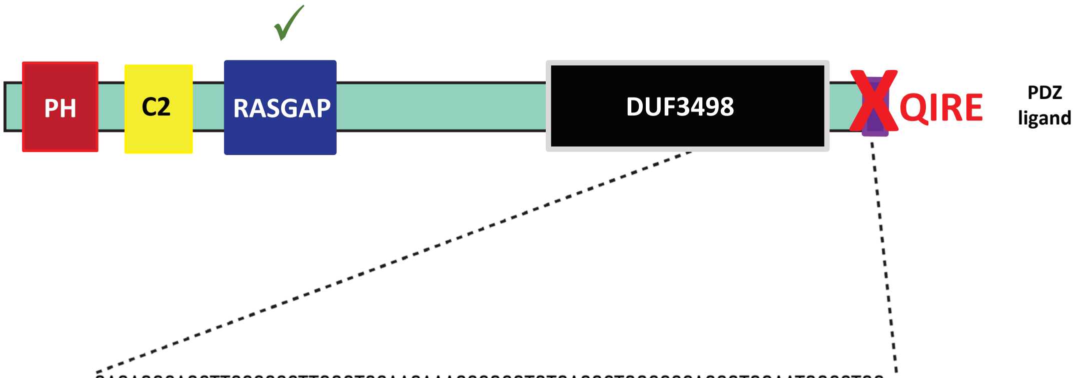


WT(03231) Genomic DNA
...GAGAGGCAGCTTCCCCCCTTGGGTCCAACAAACCCG<u>CGTGTGACGCTGGCCCCACC</u>GTGGAATGGCCTGG ccccccagccccaccaccccaccccGGCTGC...

ssODN
...CCTTTTTGGTGTCTTGCAGGAGAGGCAGCTTCCCCCC TTGGGTCCAACAAATCCGCGAGTGACGCTGGCCCCA CCGTGGAATGGCCTGGCCCCCCCAGCCCCACCACCC...

B

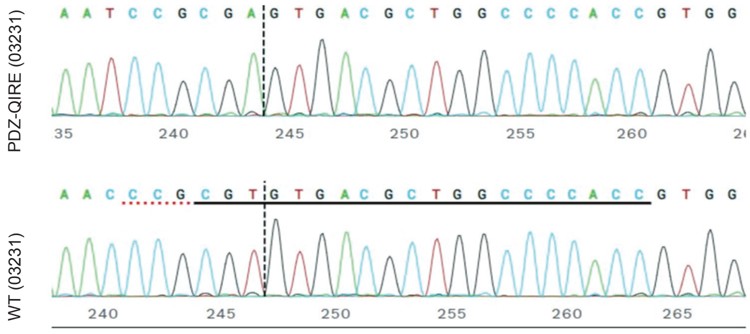


C

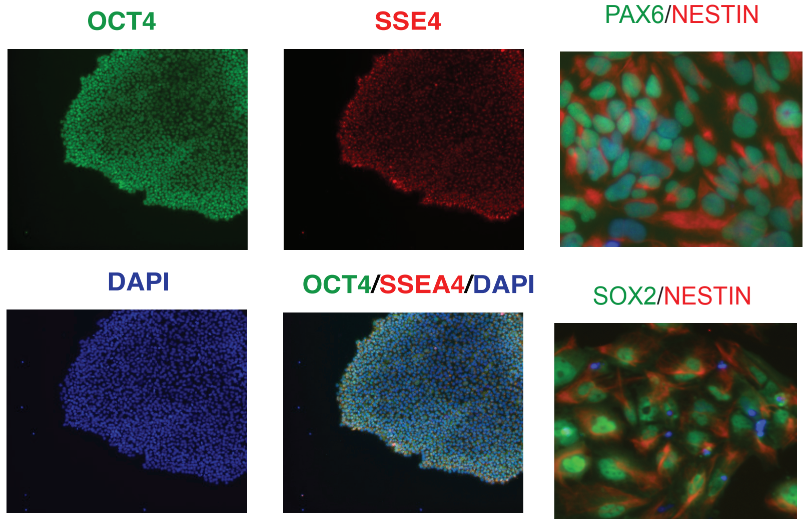


D

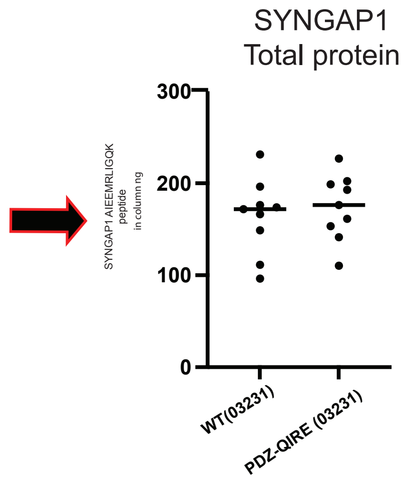
