## Supplementary material for "Disruption of the SYNGAP1 PDZ ligand motif accelerates differentiation of human iPSC-derived GABAergic neurons": Supp Fig 2

Suppl. Figure 2

### Phosphopeptides disregulated in SYNGAP1 PDZ-QIRE(03231) GABAergic iN

**Phosphopeptides upregulated (Total)**

Synaptic localization

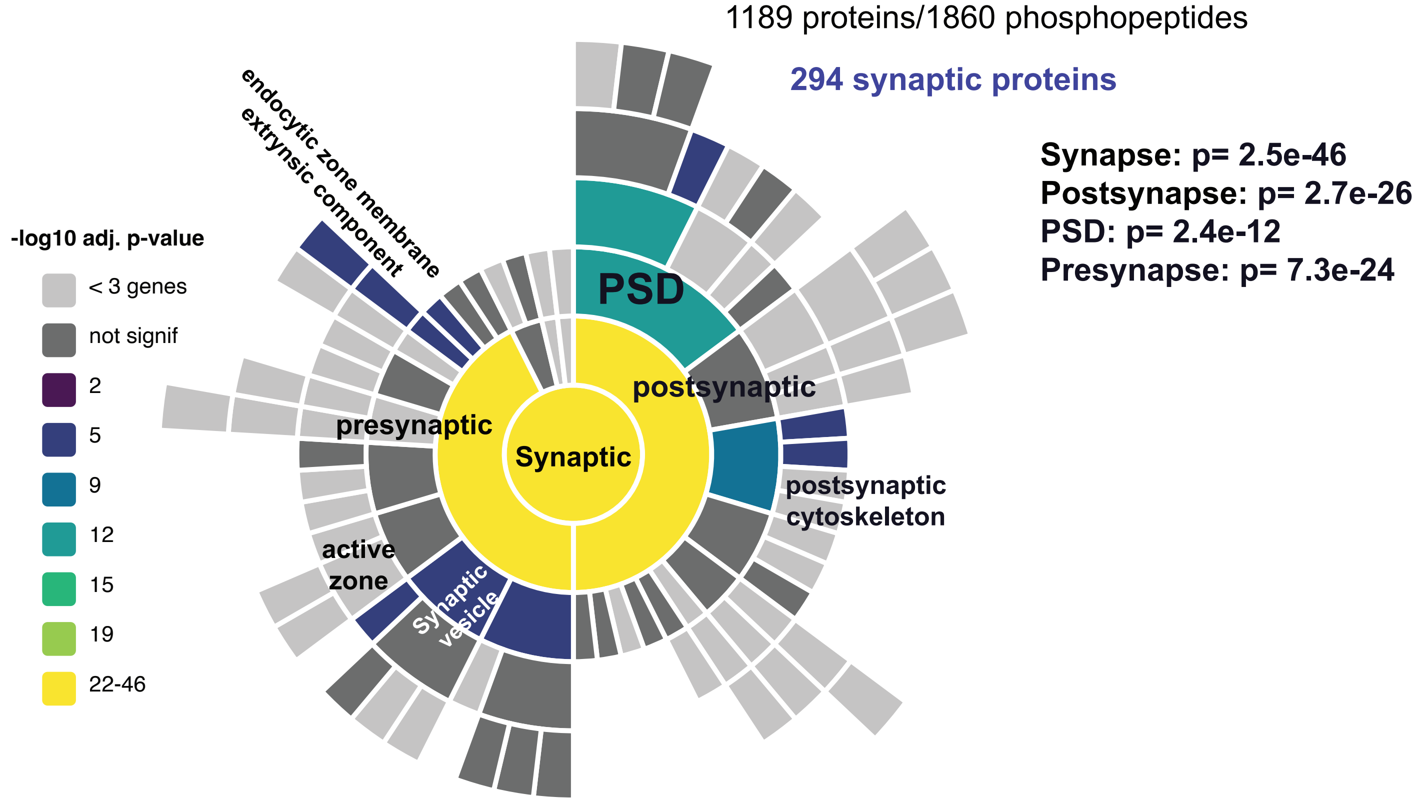


Synaptic functions

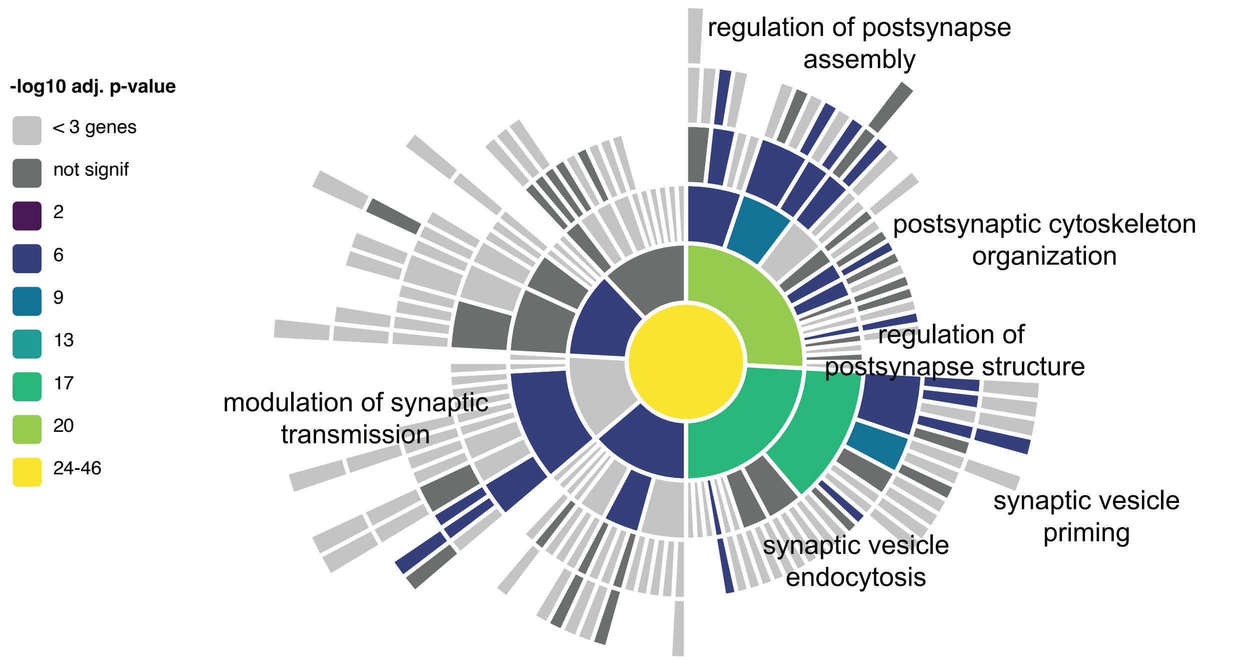
