## Supplementary material for "Disruption of the SYNGAP1 PDZ ligand motif accelerates differentiation of human iPSC-derived GABAergic neurons": Supp Fig 3

**Suppl. Figure 3**

### Phosphopeptides upregulated (no changes in total protein levels)

A

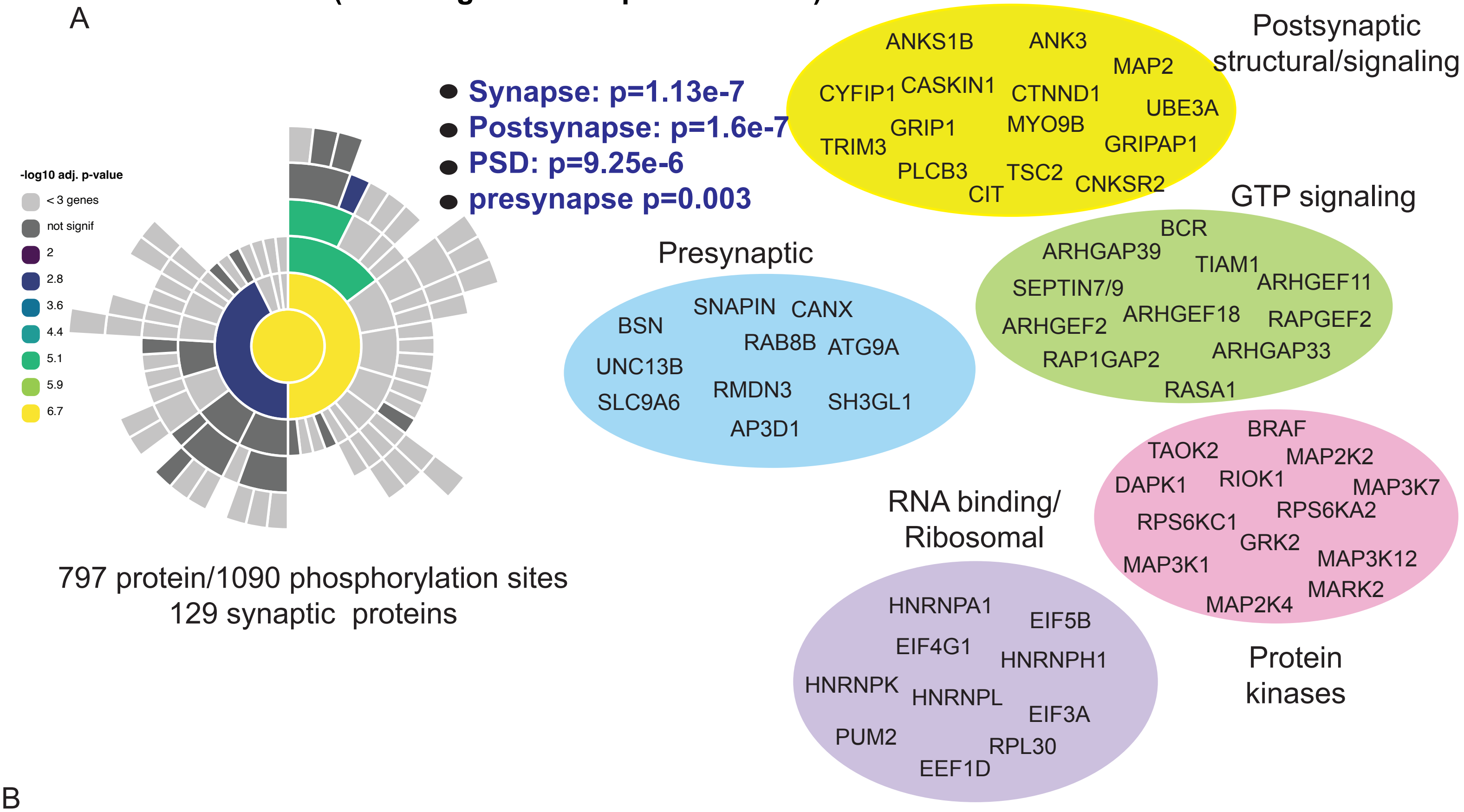


B

### Disregulated phosphopeptides: (Non-synaptic component)

#### Top canonical pathways

| Name | p-value |
|---|---|
| Processing of Capped Intron-Containing Pre-mRNA | 7.08E-21 |
| RNA Polymerase II Transcription | 3.16E-11 |
| NER (Nucleotide Excision Repair, Enhanced Pathway) | 4.80E-09 |
| Nucleotide Excision Repair | 9.27E-08 |
| Mitotic Metaphase and Anaphase | 9.34E-08 |

#### Molecular and Cellular Functions

| Name | p-value range | # Molecules |
|---|---|---|
| RNA Post-Transcriptional Modification | 2.09E-03 - 2.60E-31 | 65 |
| Gene Expression | 4.00E-05 - 7.38E-22 | 166 |
| Cellular Function and Maintenance | 2.44E-03 - 1.28E-20 | 280 |
| Cellular Development | 2.10E-03 - 1.44E-19 | 233 |
| Cellular Growth and Proliferation | 2.35E-03 - 1.44E-19 | 232 |
